# Neuronal regulated Ire1-dependent mRNA decay controls germline differentiation in *C. elegans*

**DOI:** 10.1101/2020.12.29.424718

**Authors:** Mor Levi-Ferber, Rewayd Shalash, Adrien Le Thomas, Yehuda Salzberg, Maor Shurgi, Avi Ashkenazi, Sivan Henis-Korenblit

## Abstract

Understanding the molecular events that regulate cell pluripotency versus acquisition of differentiated somatic cell fate is fundamentally important. Studies in *C. elegans* demonstrate that knockout of the germline-specific translation repressor *gld-1*, causes germ cells within tumorous gonads to form germline-derived teratoma. Previously we demonstrated that ER stress enhances this phenotype to suppress germline tumor progression (Levi-Ferber M, 2015). Here, we identify a neuronal circuit that non-autonomously suppresses germline differentiation, and show that it communicates with the gonad via the neurotransmitter serotonin to limit somatic differentiation of the tumorous germline. ER stress controls this circuit through regulated IRE-1-dependent mRNA decay of transcripts encoding the neuropeptide FLP-6. Depletion of FLP-6 disrupts the circuit’s integrity and hence its ability to prevent somatic-fate acquisition by germline tumor cells. Our findings reveal mechanistically how ER stress enhances ectopic germline differentiation, and demonstrate that RIDD can affect animal physiology by controlling a specific neuronal circuit.

## Introduction

Pluripotency is the developmental potential of a stem cell to give rise to cells of the three embryonic germ layers. Pluripotent stem cells maintain a proliferative, undifferentiated state as long as they reside within a “niche” that provides continuous pro-mitotic, anti-differentiation cues [1]. Exit from this niche is associated with a tightly-regulated switch from self-renewal to differentiation, demonstrating that the immediate micro-environment plays a central role in regulating cell pluripotency. Ultimately, the progressive restriction of cell fate is driven cell autonomously by cell-intrinsic mechanisms that regulate the expression of differentiation promoting genes, including pluripotency-regulatory transcription factors and chromatin remodeling factors, as well as translational control of gene expression [2].

The germline stem cells are the ultimate pluripotent cells, because they hold the potential to differentiate into all the cell types that comprise the embryo and the adult. Germ cells initially give rise to differentiating gametes, whose somatic fate must be repressed until fertilization. Nevertheless, at a low frequency, germ cells and gametes can generate rare germline tumors containing differentiated somatic cells that are not native to the location in which it occurs [3, 4]. These somatic germline tumors are called teratomas. In humans, the most common form of teratoma originates from oocytes that have entered, but not properly completed, meiosis [5]. The mere existence of ovarian teratomas suggests that oocyte pluripotency is normally restrained to prevent uncontrolled precocious differentiation into somatic cells.

The *C. elegans* gonad provides a well-defined model for studying how germline cells maintain pluripotency in their natural micro- and macro-environments, *i.e*. in the somatic gonad and in the whole organism [6]. The *C. elegans* germline stem cells are located in the distal end of the gonad, near the mesenchymal distal tip cell, which interacts with the germ cells and promotes germ line stemness [7]. As in humans, germ cell differentiation is initially limited to meiotic progression and gametogenesis, until the removal of somatic differentiation constraints occurs upon fertilization [8]. Nevertheless, at a low frequency (enhanced by certain genetic backgrounds), germ cells can abnormally differentiate into many kinds of somatic cells in the absence of fertilization [3, 9]. We refer to these germline cells, which lose their pluripotency and acquire a differentiated somatic cell fate prior to fertilization, as GED (germline ectopic differentiation). In *C. elegans*, GED can occur by direct conversions of germline cells to somatic cells. Such conversions have been observed upon ectopic expression of a differentiation-promoting transcription factor in the germline [10] and upon depletion of the germline-specific translation repressor *gld-1* [3, 9]. In the absence of *gld-1*, female germline stem cells initiate meiosis, but then exit pachytene and return to the mitotic cycle, yielding a tumorous germline phenotype [11]. A fraction of the mitotic germ cells forms a teratoma, which expresses somatic markers and is characterized by aberrantly large, misformed nuclei, readily detectable by DAPI staining [3, 9].

The endoplasmic-reticulum (ER) mediates correct folding of secretory proteins. ER stress increases the load of misfolded proteins in the ER. In turn, the accumulation of misfolded proteins in the ER, activates an ER-adaptive unfolded protein response (UPR). The ER-UPR consists of signaling pathways that help to restore ER homeostasis by reducing the load on the ER and degrading misfolded proteins [12]. Previously we have shown that activation of the conserved UPR sensor IRE-1 (inositol requiring enzyme-1) enhances ectopic differentiation of the tumorous germline in *gld-1*-deficient *C. elegans*, which limits the progression of the lethal germline tumor [13]. Although the major signaling mode of action of IRE-1 is to generate an active form of the ER stress-related transcription factor *xbp-1* (X-box binding protein-1), ER stress-induced GED requires the *ire-1* gene, but not its downstream target *xbp-1*. The nature of this *ire-1-*dependent *xbp-1*-independent signal remains a mystery [13]. *xbp-1*-independent outputs of *ire-1* include activation of signaling cascades by virtue of IRE-1 oligomerization [14, 15], and degradation of ER-localized mRNAs by virtue of IRE-1 ribonuclease activity—a process called regulated Ire1-dependent decay (RIDD) [16, 17]. During RIDD, select mRNAs and microRNAs are nicked by IRE1, rendering them vulnerable to rapid degradation by cytoplasmic exoribonucleases [16, 18, 19]. Thus far RIDD has been implicated in a variety of biological processes, including regulation of programmed cell death, cellular differentiation and inflammation [20–22].

In this work, we set out to explore how ER stress modulates the germ cell pluripotency/differentiation switch in the tumorous germline of *gld-1*-deficient animals. We discovered that the differentiation of the germline in response to ER stress is regulated at the organismal level, and is based on neuronal signaling. We identified a novel ER stress-sensitive neuronal circuit, which communicates with the gonad through the conserved neurotransmitter serotonin to actively repress ectopic germline differentiation. Surprisingly, it is not the stress per se that regulates germ cell fate, but rather the activation of the ER stress response sensor IRE-1, which destabilizes through RIDD the transcripts of a critical neuropeptide implicated in the GED regulatory neuronal circuit. To our knowledge this is the first example of neuronal circuit suppression by RIDD.

## Results

### Neuronal IRE-1 regulates ER stress-induced GED

The tumorous germline of *gld-1(-)* animals is prone to undergo ectopic differentiation into somatic cells, which are cleared away from the gonad by programmed cell death followed by corpse absorbance by the surrounding tissues [3, 13]. These abnormal, ectopically differentiated cells can be identified based on the expression of somatic markers and based on their relatively enlarged and irregularly shaped nuclei, which are distinct from the typical nuclei of the germline cells. The expression of somatic markers by this subgroup of cells with aberrant nuclei within the gonad of ER-stressed *gld-1(-)* animals has been previously demonstrated, confirming their somatic state (see Figure 3 - figure supplement 1 in [13]). Blocking the ability of the ectopic somatic cells to execute apoptosis increases their half-life in the gonad, facilitating their detection in nearly all of the animals, where they occupy up to 10% of the gonad [13]. Previously we have shown that upon exposure to ER stress the population of ectopically differentiated germ cells expands and occupies up to 40% of the gonad of *gld-1(-)* animals, indicating that ER stress promotes the induction of a somatic differentiated state [13]. Given the central role of the ER in secretory-protein biosynthesis, including the processing of a variety of hormones and neuropeptides, we wondered whether ER stress-induced GED is a consequence of stress within the germline itself, or the perturbation of non-autonomous signals emanating from outside the germline.

We first examined the possibility that ER stress within the germ cells controls GED. To induce ER stress primarily in the germ cells, we used mutants in the *rrf-1* gene, whose RNAi activity is compromised in most somatic tissues but whose germline RNAi activity is intact [23]. Specifically, *rrf-1* mutants were treated with a mixture of *gld-1, tfg-1* and *ced-3* RNAi. *gld-1* RNAi induces a germline tumor and sensitizes the germline for GED [3]. *tfg-1* RNAi induces ER stress by abrogating protein export from the ER [24, 25]. *ced-3* RNAi blocks apoptosis [26] and extends the half-life of the ectopic somatic cells in the gonad, facilitating their detection [13]. The efficacy of each of the RNAi treatments was individually confirmed as described in the methods section. To follow GED, the animals were stained with DAPI, and the presence of ectopically large somatic-like nuclei with irregular shape within the gonad was scored. Strikingly, even though *rrf-1* is not required for RNA interference in the germline, hardly any increase in the amount of ectopic somatic cells in the gonad was apparent upon treatment of *rrf-1* mutants with the *gld-1/tfg-1/ced-3* RNAi **(Fig. 1A)**. This was in contrast to wild-type animals with intact RNAi machinery, in which approximately 25% of each gonad was occupied by cells with large soma-like nuclei upon exposure to ER stress (**Fig. 1A**). This data suggests that ER stress in the soma, rather than specifically in the germline, drives the induction of aberrant cells within the gonad.

**Figure 1.**
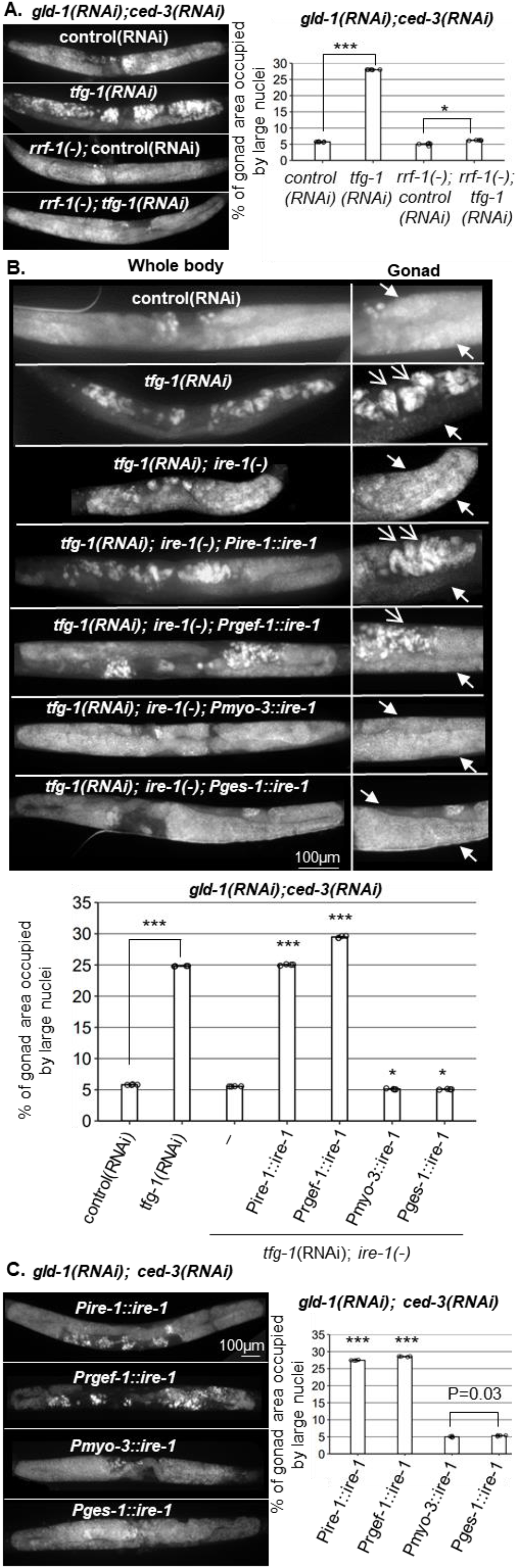
ER stress in the soma regulates GED. Percent of gonad area occupied by aberrant somatic-like cells determined by DAPI-staining of day-4 *gld-1(RNAi); ced-3(RNAi)* animals. **(A)** *tfg-1* RNAi treatment resulted in high GED levels in wild-type animals but not in *rrf-1* mutants (n= 320 gonads per genotype, N=6). **(B)** Rescue of *ire-1* expression in the soma *(Pire-1::ire-1)* and in the neurons *(Prgef-1::ire-1)* promoted GED upon *tfg-1* RNAi treatment, whereas expression of *ire-1* in the muscles *(Pmyo-3::ire-1)* and in the intestine *(Pges-1::ire-1)* did not. Full arrows indicate mitotic germ cells. Open arrows indicate aberrant nuclei (n= 210 gonads per genotype, N=4). **(C)** Overexpression of *ire-1* in the soma *(Pire-1::ire-1)* and in the neurons *(Prgef-1::ire-1)* of animals with wild-type *ire-1* resulted in high GED levels even in the absence of ER stress (n= 250 gonads per genotype, N=4). Asterisks mark one-way ANOVA followed by Tukey’s post hoc analysis of p < 0.001. Triple asterisks mark significant results resulting in at least a 2-fold change. Comparisons were made against ire-1(-) animals (B) and Pges-1::ire-1 animals (C), unless indicated otherwise.

Because IRE-1 is required for ER stress-induced GED [13], we examined the site of action of IRE-1 within the soma with respect to GED. To this end, we used *ire-1(-)* mutants, whose germline does not undergo GED in response to ER stress induced by *tfg-1* RNAi treatment [13], and restored IRE-1 expression in different somatic tissues. To restore IRE-1 expression in the entire soma but not the germline, we generated *ire-1(-)* animals expressing an extrachromosomal somatic *ire-1* transgene under its own promoter (note that extrachromosomal arrays are actively silenced in the *C. elegans* germline, but are expressed in the soma [27]). This line of experiments was carried out in animals treated with *gld-1* and *ced-3* RNAi, to facilitate GED detection. To follow GED, the animals were stained with DAPI, and the presence of abnormal nuclei within the gonad was scored. We found that expression of the somatic IRE-1 transgene was sufficient to fully restore the induction of aberrant cells within the gonad upon ER stress.

Next, we examined which somatic tissues mediate ER stress-induced GED. To this end, we restored IRE-1 expression in specific somatic tissues of *ire-1(-)* mutants by driving expression of the rescuing transgene with tissue-specific promoters. We found that expression of *ire-1* under the *rgef-1* pan-neuronal promoter was sufficient to permit ER stress-induced abnormal somatic-like nuclei within the gonad of *ire-1(-)* animals. In contrast, expression of *ire-1* under the *myo-3* muscle promoter or under the *ges-1* intestine promoter did not increase the level of abnormal nuclei in the gonads of ER-stressed animals **(Fig. 1B)**.

While IRE-1 is normally activated under ER-stress conditions, some activation of IRE-1 can be achieved merely by its over-expression [28, 29]. Hence, we asked whether increasing IRE-1 expression is sufficient for inducing GED even in the absence of direct ER stress. To this end, we overexpressed *ire-1* transgenes in various tissues of *ire-1(+); gld-1(RNAi); ced-3(RNAi)* animals and followed GED induction in the absence of an ER stress trigger. Overexpression of IRE-1 in wild-type animals, either in the soma or in the neurons gave rise to many aberrant cells within the gonad, even in the absence of ER stress **(Fig. 1C)**. In contrast, the abundance of aberrant cells in the gonad was not increased upon IRE-1 over-expression in muscles or in the intestine **(Fig. 1C)**. Since over-expression of IRE-1 is sufficient for its artificial activation independent of ER stress, this data suggests that active IRE-1 signaling in the neurons, rather than neuronal ER stress or ER dysfunction, promotes the formation of aberrant cells in the tumorous gonads of these animals.

### ER stress-induced GED is dictated by specific sensory neurons

Next, we tested whether IRE-1-induced GED is controlled by specific neurons or in a neuron-wide manner. This was examined by expressing *ire-1*-rescuing constructs driven by various neuronal promoters in *ire-1(-); gld-1(RNAi); ced-3(RNAi)* animals and scoring for restoration of ER stress-induced GED. We found that rescue of *ire-1* expression in dopaminergic or GABAergic neurons (driven by the *dat-1* and *unc-25* promoters respectively), did not induce aberrant cells in the gonads under ER stress conditions **(Fig. 2A and Figure 2 - figure supplement 1)**. In contrast, rescue of *ire-1* expression in sensory neurons or in glutamatergic neurons (driven by the *che-12* and *eat-4* promoters), led to high levels of aberrant cells in the gonads under ER stress conditions **(Figs. 2A and Figure 2 - figure supplement 1)**. Note that although the *che-12* promoter drives expression in 16 sensory neurons, only 10 of them overlap with the *eat-4* promoter **(Table S1)**. Altogether, these findings indicate that the increased generation of aberrant cells upon ER stress is dictated by specific sensory neurons, and it is not a neuron-wide phenomenon. Furthermore, the relevant neurons could be the overlapping subset of sensory and glutamatergic neurons.

**Figure 2.**
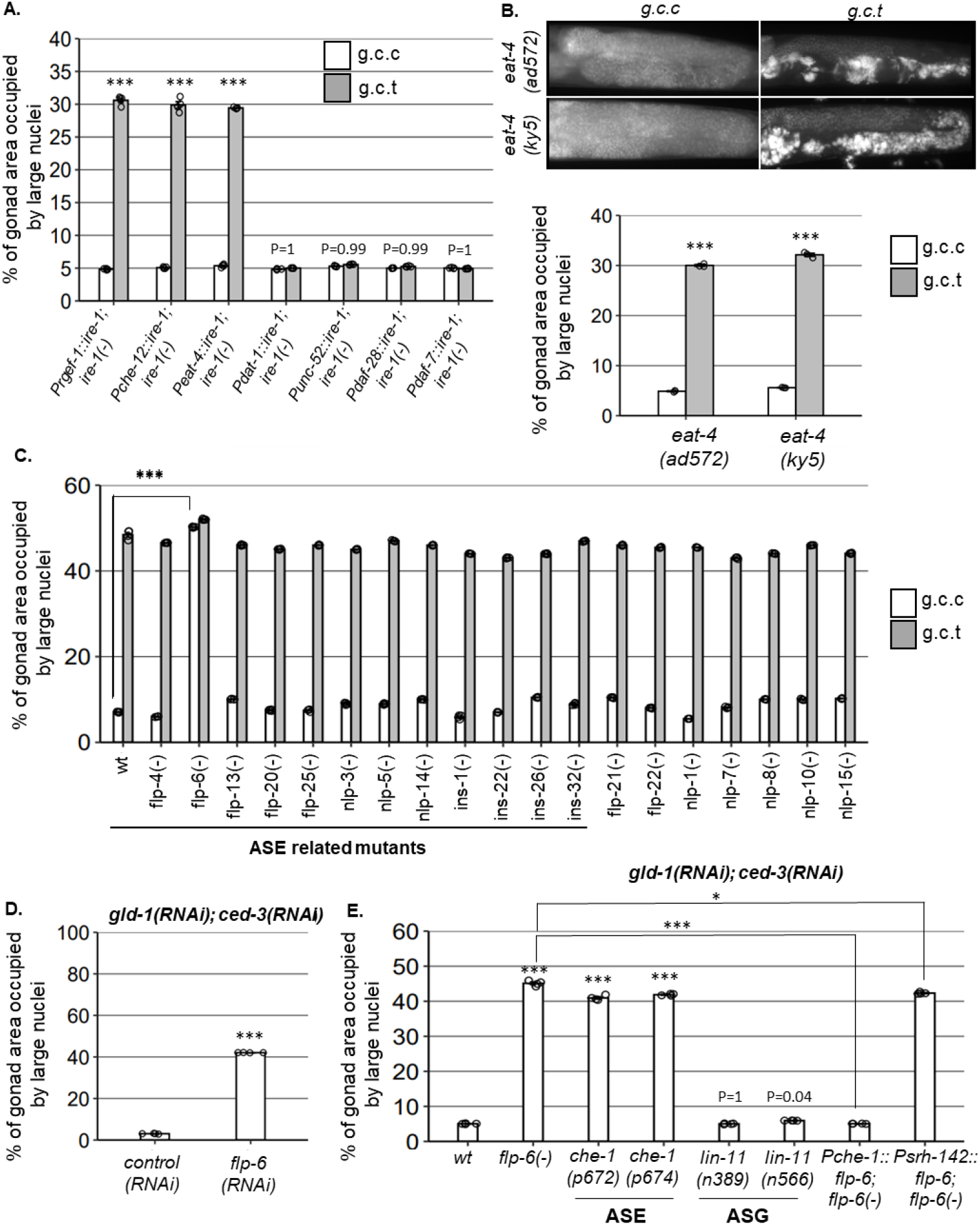
ASE-secreted FLP-6 regulates GED. Percent of gonad area occupied by ectopic somatic cells determined by DAPI-staining of day-4 animals. *g.c.c* represents treatment with *gld-1, ced-3* and control RNAi mixture. *g.c.t* represents treatment with *gld-1, ced-3* and *tfg-1*RNAi mixture. **(A)** Rescue of *ire-1* expression in all neurons *(Prgef-1::ire-1)*, in sensory neurons *(Pche-12::ire-1)* or in glutamatergic neurons *(Peat-4::ire-1)* resulted in high GED levels upon *tfg-1* RNAi treatment, whereas expression of *ire-1* in the dopaminergic *(Pdat-1::ire-1)*, GABAergic *(Punc-25::ire-1)* and *ASI/ASJ* neurons *(Pdaf-28::ire-1* and *Pdaf-7::ire-1)* did not (n= 210 gonads per genotype, N=4). Asterisks mark two-way ANOVA followed by Tukey’s post hoc analysis of p < 0.001 relative to the same animals treated with *g.c.c* RNAi. See figure 2 – figure supplement 1. **(B)** *eat-4* mutants displayed GED only upon *tfg-1* RNAi treatment (n=180 gonads per genotype, N=3). Asterisks mark two-way ANOVA followed by Tukey’s post hoc analysis of p < 0.001 relative to the same animals treated with *g.c.c* RNAi. **(C-D)** *flp-6* mutants (**C**) and *flp-6* RNAi-treated animals (**D**) displayed high GED levels in the absence of ER stress (n=23O gonads per genotype, N=4). Asterisks mark Student’s T-test of p < 0.001. **(E)** *che-1* mutants (with defective ASE) displayed high GED, whereas *lin-11* mutants (with defective ASG) did not. Rescue of *flp-6* expression in ASE *(Pche-1::flp-6)* suppressed GED in *flp-6(-)* mutants, whereas its expression in ADF *(Psrh-142::flp-6)* did not. (n=195 gonads per genotype, N=4). Asterisks mark one-way ANOVA followed by Tukey’s post hoc analysis of p < 0.001 relative to wild-type animals, unless indicated otherwise. All animals were treated with *gld-1* and *ced-3* RNAi. Triple asterisks mark significant results resulting in a 2-fold change or more.

### ASI does not regulate GED

Previously, we have shown that ASI regulates ER stress-induced germline apoptosis [24]. Furthermore, it is known that sensory information indicating favorable environmental conditions is relayed to the distal tip cell through altered TGFβ production by the ASI neurons [30]. ASI is included in the group of sensory neurons covered by the *che-12* promoter, however, it does not communicate with glutamate. Nevertheless, given its known role in regulation of germ cell proliferation and germ cell apoptosis, we examined whether it may also be involved in ER stress-induced GED. Expression of *ire-1* under the *Pdaf-28* promoter (which drives *ire-1* expression in ASI/ASJ neurons) or under *Pdaf-7* (which drives *ire-1* expression specifically in the ASI neuron) did not restore ER stress-induced GED in *gld-1* sensitized *ire-1(-)* animals **(Fig. 2A and Figure 2 - figure supplement 1)**. Thus, IRE-1 signaling in ASI is not sufficient for ER stress-induced GED, suggesting that the signaling pathways that regulate ER stress-induced GED and apoptosis are distinct.

### ASE-secreted FLP-6 regulates GED

Next, we hypothesized that the activity of sensory neurons, mediated by the release of glutamate or neuropeptides, communicates a signal that controls germline pluripotency in the gonad. One possibility is that ER stress in the relevant sensory neurons activates a signaling cascade that promotes GED in the gonad. If so, inactivation of the critical signaling molecules/cells of such a germline pluripotency inhibitory/ GED-promoting signal should prevent ER stress-induced GED. Alternatively, since ER stress conditions are incompatible with efficient signaling due to secretory defects [31], ER stress conditions that promote GED formation may do so by interfering with an existing germline pluripotency-promoting signal, rather than generating a novel GED-promoting signal. In this case, inactivation of the critical signaling molecules/cells of such a germline pluripotency promoting pathway should lead to the generation of GED, even in the absence of ER stress. To distinguish between these possibilities, we examined whether a deficiency in any of the potential neurotransmitters and neuropeptides, expressed in the 10 glutamatergic sensory neurons, is sufficient to prevent ER stress-induced GED in ER stressed animals, or sufficient to induce GED in non-stressed *gld-1(RNAi); ced-3(RNAi)* animals.

Since the main neurotransmitter synthesized in the 10 candidate sensory neurons is glutamate, we examined whether glutamate serves as a signaling molecule for GED. To this end, we assessed GED in two different glutamate transport deficient *eat-4(-)* mutants. Upon treatment with *ced-3* and *gld-1* RNAi, *eat-4(-)* mutants exhibited low levels of abnormal nuclei under non-stress conditions and high levels of abnormal nuclei under ER stress conditions **(Fig. 2B)**, similarly to wild-type animals **(Fig. 1A).** Thus, the interference with glutamate transport does not preclude the induction of GED under ER stress conditions, nor is it sufficient for induction of GED.

We next searched for candidate neuropeptides that may be involved in GED. To this end, we analyzed mutants that lack each one of the neuropeptides known to be expressed by the 10 suspected glutamatergic sensory neurons **(Table S1)**. Upon treatment with *ced-3* and *gld-1* RNAi, all of the mutants had high levels of abnormal nuclei in their gonads under ER stress conditions (**Fig. 2C**, black bars). Strikingly, one neuropeptide mutant – *flp-6(ok3056)*, exhibited high levels of abnormal nuclei under non-stress conditions as well (**Fig. 2C**, gray bars). Similar results were obtained by treatment with *flp-6* RNAi **(Fig. 2D)**. This indicates that the FLP-6 neuropeptide is a negative regulator of GED, consistent with the notion that ER stress may interfere with a fundamental germline pluripotency-promoting signal in the tumorous germline of *gld-1*-deficient animals.

FLP-6 is a secreted neuropeptide with putative hormonal activity, previously implicated in lifespan regulation [32]. It is part of a family of genes encoding FMRF amide-related peptides (FaRPs), which regulate many aspects of behavior [33, 34]. FLP-6 is produced in six neurons (ASE, ASG, AFD, ADF, I1, PVT) [35]. AFD-derived FLP-6 has been implicated in temperature-related longevity regulation [32]. Nevertheless, only ASE and ASG are included in the group of sensory neurons that overlap with the *che-12* and *eat-4* promoters’ expression pattern. Therefore, we further assessed ASE and ASG involvement in GED.

Since *flp-6* depletion was sufficient to induce GED in the absence of ER stress, we hypothesized that depletion of its producing cells will have a similar effect. To this end, we genetically inactivated the ASE and ASG cells and examined if this was sufficient to induce GED in *gld-1* and *ced-3 RNAi*-treated animals. We examined two *che-1* mutants, which lack a critical transcription factor involved in ASE fate specification, and thus are ASE deficient [36]. We observed high levels of GED in ASE(-) *che-1* mutants compared to wild-type animals in the absence of ER stress, resulting in similar GED levels as in *flp-6(-)* mutants **(Fig. 2E)**. In contrast, two *lin-11* mutants, which lack a critical transcription factor involved in ASG fate specification and are thus ASG defective [37], exhibited low GED levels similar to wild-type animals **(Fig. 2E)**. These observations suggest that the ASE sensory neuron actively prevents GED in *gld-1(RNAi); ced-3(RNAi)* animals.

Taking together the requirement of both ASE and *flp-6* for GED prevention, we hypothesized that ASE-produced FLP-6 suppresses GED in *gld-1(RNAi); ced-3(RNAi)* animals. To examine this, we generated *flp-6(-)* animals expressing a *flp-6* rescuing construct under the ASE specific promoter *Pche-1*. Unlike *flp-6(-); gld-1(RNAi); ced-3(RNAi)* mutants, which had high levels of abnormal nuclei in their gonads, transgenic animals expressing ASE-produced FLP-6 exhibited low levels of abnormal nuclei in their gonads, similarly to wild-type animals **(Fig. 2E).**

Finally, since FLP-6 is a secreted neuropeptide, we wondered whether its secretion by other FLP-6-producing cells may be sufficient for the suppression of GED in *gld-1(-); ced-3(-)* animals. Hence, we examined the levels of GED in *flp-6(-); gld-1(RNAi); ced-3(RNAi)* mutants upon restoration of *flp-6* expression in ADF, one of the six known FLP-6-prodcing neurons. In contrast to the animals expressing *flp-6* specifically from the ASE neurons, *flp-6(-)* animals expressing *flp-6* rescuing construct under an ADF specific promoter *(Psrh-142)* exhibited high GED levels similarly to *flp-6(-)* animals **(Fig. 2E)**. These results suggest that ASE-produced FLP-6 is a critical signaling moiety in GED regulation in *gld-1(RNAi); ced-3(RNAi)* animals. Furthermore, the fact that FLP-6 production by different sensory neurons leads to distinct phenotypes suggests that FLP-6 may act locally, through synaptic transmission, rather than distally, as a hormone distributed throughout the body cavity of the organism.

### *flp-6* deficiency induces GED independently of ER stress or *ire-1*

Since ER stress promotes GED, we checked whether *flp-6* deficiency is associated with ER stress. First, we examined the possibility that *flp-6* deficiency generates ER stress. To this end, we measured the levels of the IRE-1/UPR reporter *Phsp-4::gfp*, whose transcription is increased under conditions of ER stress. We found that the fluorescence level of the *Phsp-4::gfp* reporter did not increase in animals which lack *flp-6*. Specifically, the level of the IRE-1/UPR reporter was comparable to that of animals with intact *flp-6* when measured throughout the body of the animals **(P=0.49, Fig. 3A).** When the level of the IRE-1/UPR reporter was measured specifically in the ASE cell body, a slight decrease was observed **(P=0.0.0056, Figure 3 – figure supplement 1).** This data indicates that IRE-1 is not activated in *flp-6-* deficient animals. To verify the notion that *flp-6* deficiency-induced GED is not the result of ER stress, we examined if *flp-6* deficiency-induced GED is mediated by *ire-1*, similarly to ER stress-induced GED. We found that *flp-6(-); ire-1(-)* double mutants treated with *gld-1* and *ced-3* RNAi exhibited high levels of abnormal nuclei in their gonads under non-stress conditions, similarly to *flp-6(-); gld-1(RNAi); ced-3(RNAi)* animals **(Fig. 3B)**. Together, these results indicate that *flp-6* deficiency can induce GED independently of ER stress or *ire-1*.

**Figure 3.**
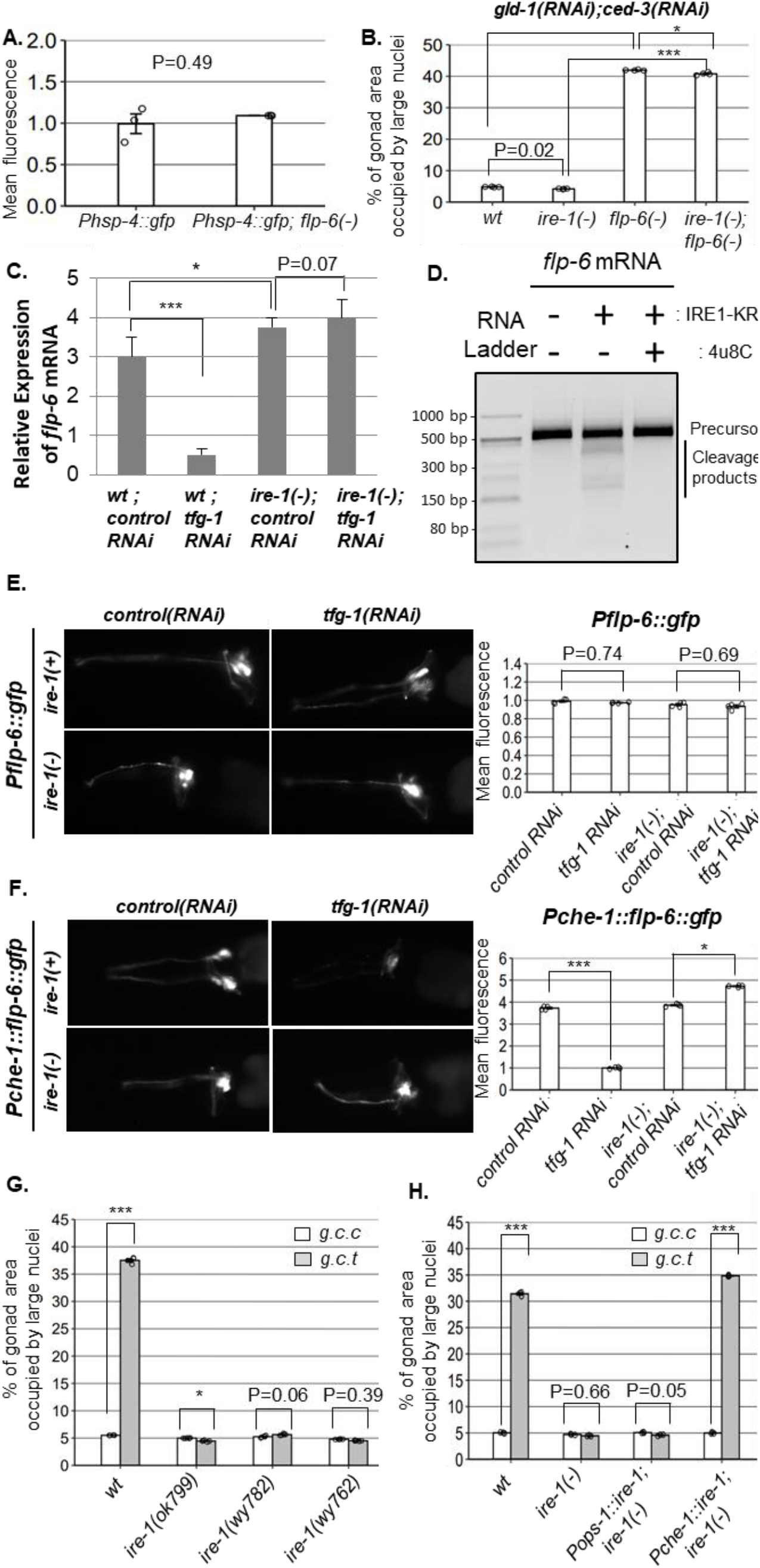
IRE-1 acts upstream of *flp-6* and controls its transcript stability. **(A)** The fluorescence levels of the *Phsp-4::gfp* IRE-1 reporter were comparable in *flp-6(ok3056)* and wild-type *flp-6(+)* day-1 animals (Student’s T-test P=0.49, n=130 gonads per genotype, N=3). See figure 3 – figure supplement 1 for ASE-specific *Phsp-4::gfp* signal. **(B)** *flp-6 ok3056* mutation increased GED independently of *ire-1*, upon treatment with *gld-1 and ced-3* RNAi (n= 200 gonads per genotype, N=4). The *ire-1 ok799* deletion allele mutant was examined. Asterisks mark one-way ANOVA followed by Tukey’s post hoc analysis of p < 0.001 as indicated. GED assessed by DAPI staining of Day-4 animals. **(C)** *flp-6* transcript levels were assessed by qRT-PCR and normalized to actin transcript levels. ER stress induced by *tfg-1* RNAi reduced *flp-6* transcript levels in wild-type animals but not in *ire-1* mutants. *flp-6* transcript levels were increased in *ire-1(-)* animals compared to wild-type animals. Asterisks indicate Student’s t-test P values p < 0.01. **(D)** Purified recombinant human IRE-1α comprising the kinase and RNase domains (IRE1-KR) was incubated with in vitro-transcribed *flp-6* RNA fragment in the presence of vehicle or 4μ8c (5 μM). Reactions were resolved on 3% agarose gel. Cleavage products of the *flp-6* RNA fragment were observed upon its incubation with IRE1-KR, but not in the presence of the specific IRE-1 ribonuclease inhibitor 4μ8C. See Figure 3 – figure supplement 3 for predicted hairpin structure of *flp-6* RNA. **(E-F)** *tfg-1* RNAi treatment did not change the fluorescence levels of the *Pflp-6::gfp* transcriptional reporter, whereas it decreased the fluorescence of the *flp-6::gfp* translation reporter driven by the heterologous ASE-specific *che-1* promoter in an *ire-1* dependent manner (n=205 gonads per genotype, N=4). P values of one-way ANOVA followed by Tukey’s post hoc analysis are indicated. See figure 3 – figure supplement 2 for *Pche-1::mcherry* signal. **(G)** Treatment with a mixture of *gld-1; ced-3; tfg-1* RNAi *(g.c.t)* failed to induce GED in the absence of functional *ire-1* (n= 210 gonads per genotype, N=5). *ok799* is an *ire-1* deletion mutation. *wy762* is an IRE-1 endoribonuclease missense mutation. *wy782* is an IRE-1 kinase missense mutation. Asterisks mark two-way ANOVA followed by Tukey’s post hoc analysis of p < 0.01 relative to the same animals treated with *gld-1; ced-3; control* RNAi *(g.c.c)*. **(H)** Rescue of *ire-1* expression in the ASE neuron *(Pche-1::ire-1)* restored ER stress-induced GED upon treatment with a mixture of *gld-1; ced-3; tfg-1* RNAi (*g.c.t*), whereas expression of *ire-1* in the ASG neuron *(Pops-1::ire-1)* did not (n= 110 gonads per genotype, N=3). g.c.c indicates treatment with a mixture of *gld-1; ced-3; control* RNAi. Asterisks mark two-way ANOVA followed by Tukey’s post hoc analysis of p < 0.001 relative to the same animals treated with *gld-1; ced-3; control* RNAi (*g.c.c*). In all panels - triple asterisks mark significant results resulting in a 2-fold change or more.

### IRE-1 acts upstream of FLP-6 and controls its transcript abundance via RIDD

GED induction by ER stress in *gld-1(-)* animals is *ire-1-*dependent but *xbp-1*-independent [13]. A key *xbp-1*-independent output of *ire-1* is RIDD [16], which degrades transcripts based on their proximity to the ER and the presence of a specific hairpin motif [38]. Although RIDD is conserved from yeast to human, surprisingly, no RIDD target has been demonstrated thus far in *C. elegans*. We hypothesized that ER stress activates *ire-1*, which in turn degrades *flp-6* transcripts encoding a protein that normally prevents GED. To test this hypothesis, we measured the mRNA levels of *flp-6* under non-stress and under ER stress conditions *(i.e tfg-1* RNAi treatment), in *ire-1(+)* and *ire-1(-)* animals, using qRT-PCR. We found that *flp-6* transcript levels were reduced under ER stress conditions in an *ire-1* dependent manner **(Fig. 3C)**. Furthermore, *flp-6* transcript levels were higher in *ire-1(-)* animals compared to wild-type animals under stress and non-stress conditions **(Fig. 3C, P<0.01).** Thus, *flp-6* transcript levels are regulated by ER stress in an *ire-1*-dependent manner.

To examine whether these ER stress-related changes in *flp-6* transcript levels are due to transcriptional or post-transcriptional regulation, we analyzed the expression of a *Pflp-6::gfp* transcriptional reporter. Analysis of the *Pflp-6::gfp* transcriptional reporter strain showed no significant change in the transcriptional activity through this promoter upon ER stress or upon *ire-1* depletion **(Fig. 3E)**. Furthermore, analysis of a *flp-6::gfp* translational reporter, under the control of the heterologous ASE-specific *che-1* promoter, did show that FLP-6 protein levels were reduced by ER stress in wild-type animals, but not in *ire-1(-)* animals **(Fig. 3F).** The reduced levels of FLP-6 protein reporter in ER-stressed wild-type animals was not due to reduced activity of the driving promoter, whose activity was unaltered by ER stress (**Fig. 3 – supplement figure 2**). This data indicates that *ire-1* decreases FLP-6 levels post-transcriptionally. Taken together with the fact that *flp-6* transcript levels are regulated by ER stress in an *ire-1*-dependent manner, these findings suggest that *flp-6* transcript is a RIDD substrate.

RIDD substrates typically harbor an *xbp-1-like* stem-loop structure comprising a seven nucleotide loop (-C-X-G-C-X-X-X-) with three conserved residues and a stem of at least four base pairs [39–41]. Within the spliced *flp-6* sequence we identified one potential *xbp-1-like* stem-loop structure **(Fig. 3 - supplement figure 3)**. Although the identified structure contains only 6 (rather than the typical 7) nucleotides in the loop, it has all the conserved residues (-C-x-G-C-X-X-) followed by a relatively strong stem structure. To test whether this RNA molecule could indeed be cleaved directly by the IRE-1 endoribonuclease, we incubated a 513bp RNA segment harboring this stem loop with a phosphorylated form of purified human IRE-1 kinase-RNase domain (IRE-1 KR-3P) [22]. Indeed, incubation of the *flp-6* stem-loop RNA fragment with purified recombinant IRE-1 KR-3P generated clearly detectable cleavage products **(Fig. 3D)**. Furthermore, the cleavage was blocked by the IRE-1-specific RNAse inhibitor 4μ8c [42]. This data indicates that the *flp-6* RNA can be directly cleaved by IRE-1, thus supporting the possibility that *flp-6* is a direct RIDD substrate.

If *flp-6* levels are down-regulated by IRE-1’s RIDD activity, and given that it is ASE-produced FLP-6 that is critical for GED prevention, we predicted that the ribonuclease activity of IRE-1 would be essential for ER stress-induced GED and that IRE-1 expression specifically in the ASE sensory neuron should be sufficient for restoration of ER stress-induced GED in otherwise *ire-1*-deficient mutants. Indeed, analysis of a kinase-dead *(wy782)* and a ribonuclease-dead *(wy762) ire-1* mutant [43] indicated that both the kinase and ribonuclease activities of IRE-1 were required for ER-stress induction of GED **(Fig. 3G).** Note that the kinase activity is required for the ribonuclease activity, which directly executes RIDD [44]. Furthermore, we individually restored *ire-1* expression in ASE or ASG (two FLP-6-producing glutamatergic sensory neurons). As expected, *Pche-1::ire-1; ire-1(-)* transgenic animals, that express *ire-1* specifically in the ASE neuron, had high levels of abnormal nuclei in their gonads upon exposure to ER stress. In contrast, *Pops-1::ire-1; ire-1(-)* transgenic animals, which express *ire-1* in the ASG and ADL sensory neurons, did not have high levels of abnormal nuclei in their gonads upon exposure to ER stress **(Fig. 3H)**. Altogether, these findings further support the conclusion that IRE-1 limits FLP-6 protein levels in ASE neuron by destabilizing *flp-6* transcripts via RIDD.

### AIY prevents GED by Acetylcholine (ACh) signaling

ASE is one of the head amphid neurons, and it does not synapse with the gonad. Hence, we examined the putative involvement of four interneurons (AIA, AIB, AIY, RIA) known to synapse with ASE [45] in the GED process. Our data support the hypothesis that ER stress conditions promoting GED formation interfere with an existing germline pluripotency-inducing signal. Accordingly, inactivation/interference of the critical signaling cells should also lead to the generation of GED, even in the absence of ER stress, as seen in ASE(-)/*flp-6*(-) animals. Hence, we used a system for inducible silencing of specific neurons, based on transgenic nematodes engineered to produce the inhibitory *Drosophila* histamine-gated chloride channel (HisCl1) in each of the 4 suspected interneurons (AIA, AIB, AIY, RIA) [46]. GED was assessed in each of the strains, with or without histamine, upon treatment with *gld-1* and *ced-3* RNAi. Whereas histamine-induced inactivation of AIA, AIB and RIA resulted in low levels of abnormal nuclei in the gonads, animals in which the AIY neuron was inactivated displayed high levels of abnormal nuclei in their gonads **(Fig. 4A)**. This result implicates AIY in GED regulation.

**Figure 4.**
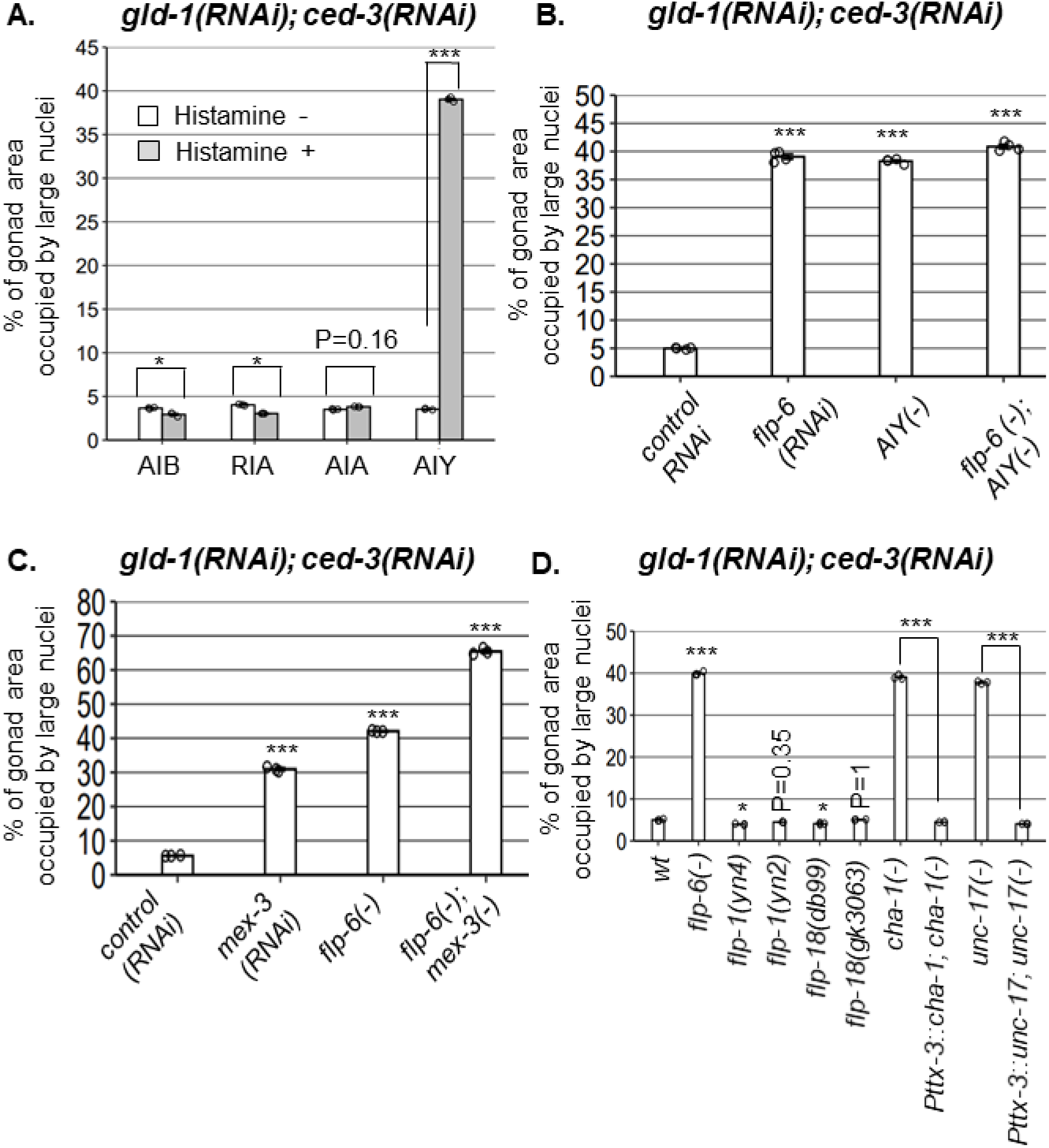
AIY produces Acetylcholine (ACh) to prevent GED. Percent of gonad area occupied by ectopic somatic cells determined by DAPI-staining of day-4 animals treated with a mixture of *gld-1* and *ced-3* RNAi. **(A)** Four histamine-inducible interneurons silencing mutants (AIA, AIB, AIY, RIA) were examined (n=170 gonads per genotype, N=3). While AIA, AIB and RIA histamine-treated animals exhibited low levels of GED, AIY histamine-treated animals had high GED levels. Asterisks mark two-way ANOVA followed by Tukey’s post hoc analysis of p < 0.001 relative to the same animals without histamine treatment. **(B)** *flp-6* RNAi did not additively increase GED levels in AIY histamine-treated animals (n=180 gonads per genotype, N=4). P values were determined by one-way ANOVA followed by Tukey’s post hoc analysis. All strains were grown in the presence of histamine. **(C)** Animals deficient in both *flp-6(ok3056)* and *mex-3* additively increased GED (n=215 gonads per genotype, N=3). Asterisks mark oneway ANOVA followed by Tukey’s post hoc analysis of p < 0.001 relative to animals treated with the *gld-1, ced-3, control* RNAi mix. **(D)** *cha-1(p1152)* and *unc-17(e113)* mutants had high GED levels upon *gld-1* and *ced-3* RNAi treatment (n= 190 gonads per genotype, N=3). GED levels were suppressed upon expression of the corresponding transgenes in the AIY neuron (*Pttx-3::cha-1 and Pttx-3::unc-17*). Asterisks mark one-way ANOVA followed by Tukey’s post hoc analysis of p < 0.001 relative to wild-type animals unless indicated otherwise. In all panels - triple asterisks mark significant results resulting in a 2-fold change or more.

In order to see if ASE-produced *flp-6* and AIY act in a common pathway to regulate GED, we compared the levels of abnormal nuclei between *flp-6* and AIY defective animals in a *gld-1(-); ced-3(-)* genetic background. We found that each perturbation, individually or in combination, resulted in occupation of ~40% of the gonad with abnormal nuclei **(Fig. 4B)**. In contrast, treatment of *flp-6* mutants with *mex-3* RNAi (previously reported to increase GED in *gld-1(-)* animals [3]) did increase the occupancy of the gonad with GED beyond 40% in an additive manner (**Fig. 4C**), demonstrating that the analysis of GED levels in this system is not limited by saturation. Together, these results suggest that the AIY interneuron and *flp-6* act in a common pathway to control GED, whereas *mex-3* deficiency promotes GED via an independent pathway.

We next investigated how the signal is relayed from AIY to the gonad. AIY communicates mainly with *flp-1, flp-18* or acetylcholine (ACh). However, neither deficiencies in *flp-1* nor in *flp-18* resulted in high levels of abnormal nuclei in the gonads **(Fig. 4D)**, suggesting that these neuropeptides are not individually necessary for GED. Next, we examined ACh involvement in this signaling pathway, using mutations that partially perturb ACh synthesis. Upon treatment with *gld-1* and *ced-3* RNAi, we found high levels of abnormal nuclei in the gonads of *cha-1(p1152)* mutants, in which acetylcholine synthesis is defective [47], as well as in *unc-17(e113)* mutants, in which synaptic vesicle acetylcholine transport is defective [48] **(Fig. 4D)**. Furthermore, expression of a rescuing construct of *cha-1* or *unc-17* under the AIY specific *Pttx-3^promB^* promoter [49] led to low levels of abnormal cells in the gonads, similarly to the wild-type level **(Fig. 4D)**. Altogether, these findings implicate ACh signaling by AIY in the regulation of GED.

### HSN produces serotonin to prevent GED

How is the signaling cascade transmitted downstream to the gonad? AIY communicates via synapses with a few sensory neurons (AWA, AWC, ADF) and a few interneurons (RIA, RIB, AIZ). In addition, AIY synapses with the HSN-R motor neuron, which innervates the vulva muscles [45]. Based on its anatomical location near the vulva, HSN was the strongest candidate for further examination. Indeed, high levels of abnormal nuclei were identified upon treatment with *gld-1* and *ced-3* RNAi in two HSN deficient mutants (*her-1* and *sem-4*) as well as in three HSN migration defective mutants **(Fig. 5A)**. In addition, an epistasis analysis between *flp-6* and *sem-4* or *her-1* HSN deficient animals showed no additive effect on the levels of abnormal nuclei in the gonad **(Fig. 5A)**. Together, these results suggest that HSN and *flp-6* act in a common pathway to regulate GED.

**Figure 5.**
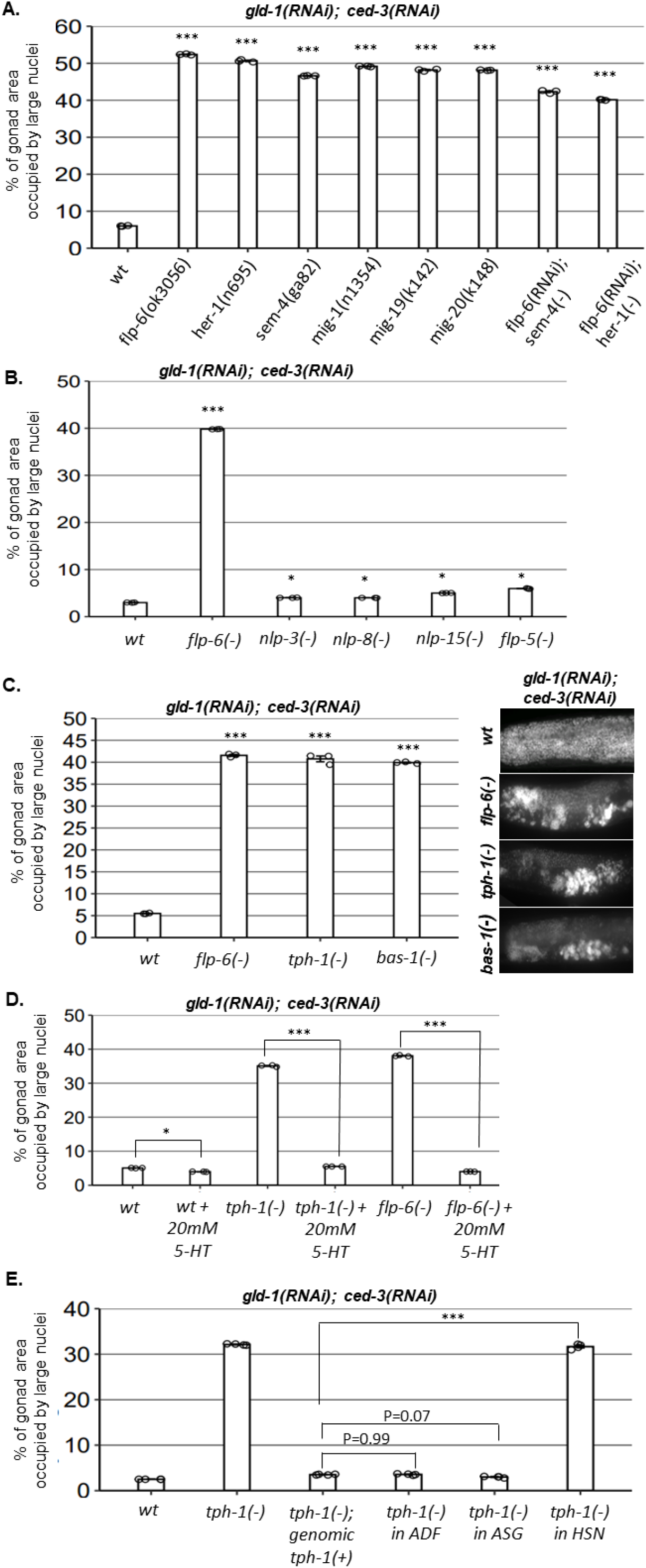
HSN produces serotonin to prevent GED. Gonad area occupied by ectopic somatic cells determined by DAPI-staining of day-4 animals treated with a mixture of *gld-1* and *ced-3* RNAi. **(A)** All HSN deficiency and migration mutants had high GED levels. *flp-6* RNAi did not further increase GED levels in HSN-defective animals (n= 175 gonads per genotype, N=3). **(B)** HSN-related neuropeptide mutants had low levels of GED, similar to wild-type animals (n= 200 gonads per genotype, N=3). **(C)** Both *tph-1(mg280) and bas-1(tm351)* serotonin mutants had high GED levels, similarly to *flp-6* mutants (n=185 gonads per genotype, N=3). See Figure 5 – supplement figure 1 for the analysis of serotonin receptor mutants. **(D)** Treatment with 20mM serotonin decreased GED levels of *tph-1(mg280)* and *flp-6(ok3056)* mutants (n= 165 gonads per genotype, N=3). Asterisks mark one-way ANOVA followed by Tukey’s post hoc analysis of p < 0.001 relative to the same animals without serotonin supplementation. **(E)** *tph-1* deficiency in HSN, but not in other serotonin-producing neurons, resulted in high GED levels (n=195 gonads per genotype, N=4). In A-E, asterisks mark one-way ANOVA followed by Tukey’s post hoc analysis of p < 0.001 relative to wild-type animals treated *gld-1* and *ced-3* RNAi, unless indicated otherwise. In all panels - triple asterisks mark significant results resulting in a 2-fold change or more.

How does HSN signal to the gonad? We examined the involvement of four HSN-produced neuropeptides (*nlp-3, nlp-8, nlp-15, flp-5*) and the neurotransmitter serotonin. None of the mutants of the HSN-related neuropeptides induced high levels of GED upon treatment with *gld-1; ced-3* RNAi **(Fig. 5B)**. In contrast, two serotonin defective mutants (*tph-1* and *bas-1*) did exhibit high levels of abnormal nuclei in their gonads, similarly to *flp-6(-)* **(Fig. 5C)**. This indicates that serotonin production is required for suppression of GED in *gld-1(-); ced-3(-)* animals, and that in its absence, GED levels increase.

There are four genes encoding major serotonin receptors in *C. elegans*. Hence, we examined whether a deficiency in any of these receptors results in GED formation in *gld-1(-); ced-3(-)* animals, similarly to animals deficient in serotonin production. However, we found that GED levels in mutants lacking each of the major serotonin receptors individually *(ser-1(-), ser-4(-), ser-7(-), mod-1(-)* in *gld-1(RNAi); ced-3(RNAi)* background) remained low **(Fig. 5 – supplement figure 1)**. This implies that in this case, serotonin affects germ cell fate by another serotonin receptor, or that it is redundantly recognized by more than one of these receptors.

After establishing that serotonin production is necessary for GED suppression, we next examined whether serotonin can suppress GED formation in *gld-1(RNAi); ced-3(RNAi)* background. To this end, we examined how externally supplied serotonin affected GED in wild-type, *flp-6(-)* and *tph-1(-)* animals. As expected, GED levels in serotonin-deficient *tph-1* mutants treated with *gld-1* and *ced-3 RNAi* was suppressed by the supplementation with 20mM serotonin **(Fig. 5D)**. Furthermore, treatment with 20mM serotonin also reduced the levels of abnormal nuclei in the gonads of *flp-6(-)* mutants treated with *gld-1* and *ced-3 RNAi* down to wild type level (38% and 5% respectively, **Fig. 5D)**. Thus, high levels of serotonin are sufficient to suppress GED induced by *flp-6* deficiency in *gld-1(-); ced-3(-)* animals.

Serotonin is produced by the ADF, ASG and HSN neurons. The fact that HSN defective *gld-1(-); ced-3(-)* animals have high levels of GED suggested that serotonin production by HSN may be critical for GED suppression. To verify that serotonin production specifically from HSN regulates GED, we measured GED levels upon *gld-1; ced-3* RNAi treatment in strains expressing Cre-Lox specific knockout of *tph-1* [50]. Inhibition of serotonin production in either ADF or in ASG resulted in low levels of abnormal nuclei in the gonads, similarly to those of the genomic rescue of *tph-1* **(Fig. 5E)**. In contrast, inhibition of serotonin production in HSN led to high levels of abnormal nuclei in the gonads, similar to those observed in *tph-1(mg280)* mutants **(Fig. 5E).** Altogether, our results suggest that HSN determines GED levels in the tumorous germline of *C. elegans* through serotonin signaling.

### Perturbation of the ASE-AIY-HSN-gonad circuit induces GED

Thus far, our findings demonstrate that the ASE, AIY and HSN neurons are required individually to prevent the accumulation of cells with abnormal nuclei in the tumorous gonad of *gld-1*-deficient animals. Previously we have shown that ER stress also results in the accumulation of cells with abnormal nuclei in the tumorous gonad of *gld-1*-deficient animals, and that these cells are ectopically differentiated somatic cells that arise from the tumorous germline [13]. Our new findings suggest that ER stress affects the differentiation state of the tumorous germline by compromising the pluripotency-protective ASE-AIY-HSN-germline circuit. Thus, the abnormal nuclei that arise and accumulate in the tumorous gonads upon perturbations within the ASE-AIY-HSN-germline circuit are likely to be the nuclei of differentiated somatic cells as well. To test this directly, we made use of a *Punc-119::gfp* panneuronal reporter, introduced into ASE-deficient *che-1* mutants, HSN-deficient *sem-4* mutants, or *flp-6/cha-1/tph-1* mutants with defective *flp-6/*acetyl-choline/serotonin signaling, all of which have been implicated in the circuit identified herein. The *Punc-119::gfp* marker is normally expressed in the nerve cells of animals, and is apparent from embryo to adult. However, upon GED in tumorous gonads, this marker is also observed in germ cells which acquired neuronal fate within the gonad [13]. Consistent with the phenomenon of GED, we found expression of the neuronal reporter within the tumorous gonads of animals with a perturbed ASE-AIY-HSN-germline but not in animals with an intact circuit (**Fig. 6A**). Taken together with the *C. elegans* connectome and epistasis analysis, we conclude that these neurons establish a neuronal circuit whose integrity prevents ectopic differentiation of the germline into somatic cells in the tumorous gonad.

**Figure 6.**
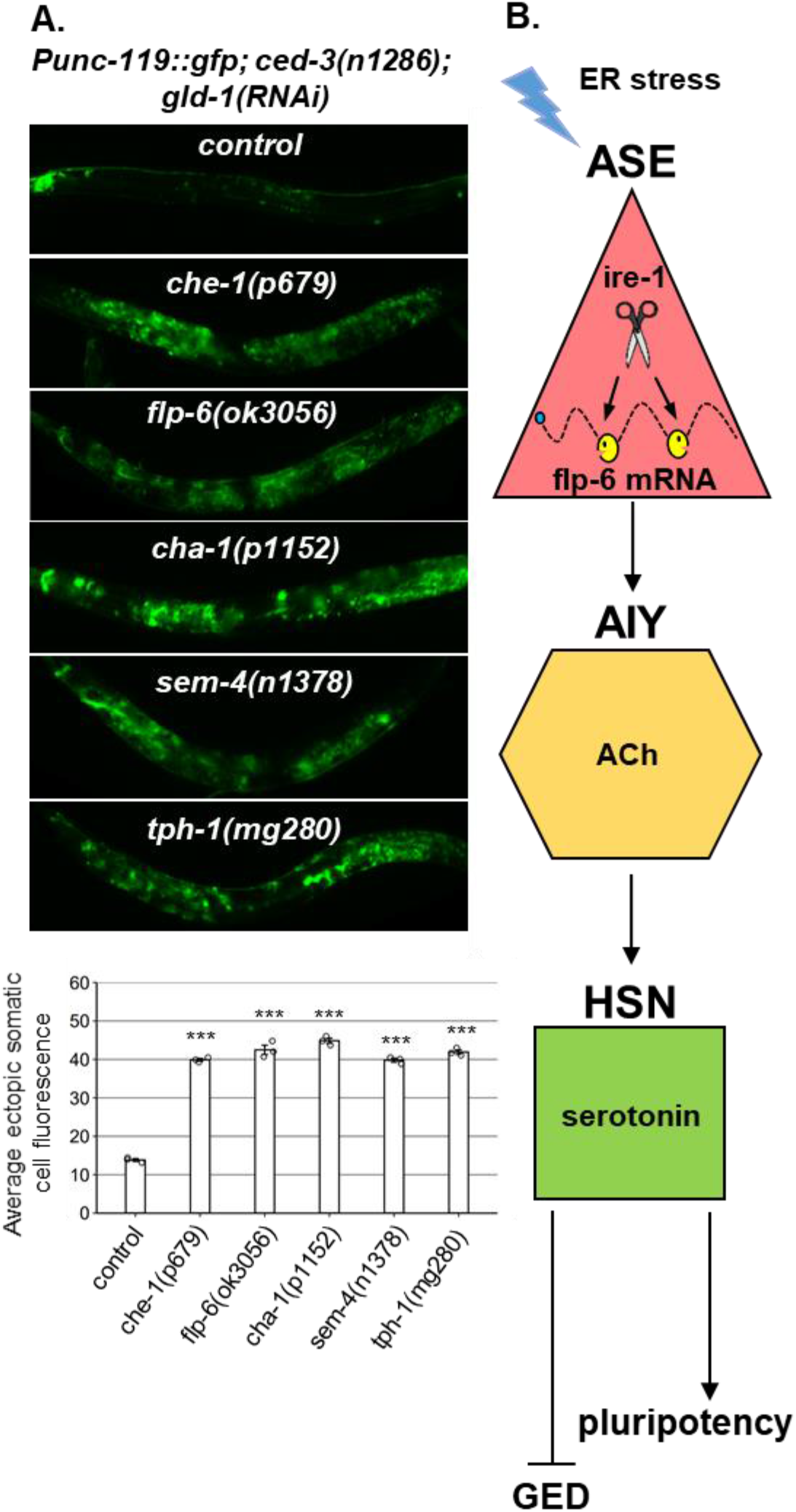
A neuronal circuit actively prevents GED. **(A)** Summarizing model of ASE-AIY-HSN-germline circuit which actively maintains germline pluripotency and suppresses ectopic germline differentiation in *gld-1* tumorous animals. Implicated neurons and signaling molecules are indicated. ER stress suppresses this circuit by RIDD-mediated destabilization of the *flp-6* transcript. This releases the inhibition from the tumorous germ cells which acquire somatic fate by default. **(B)** Abnormal expression of the neuronal reporter *Punc-119::gfp* within the tumorous gonads of animals with a perturbed ASE-AIY-HSN-germline circuit, but not in control animals with an intact circuit. *che-1* and *sem-4* mutants are ASE and HSN-deficient respectively. *flp-6* mutants are deficient in the FLP-6 neuropeptide. *cha-1* and *tph-1* mutants have compromised Ach and serotonin production. Asterisks mark one-way ANOVA followed by Tukey’s post hoc analysis of p < 0.001 relative to control *ced-3(-)* animals treated with *gld-1* RNAi. Triple asterisks mark significant results resulting in a 2-fold change or more.

## Discussion

Previous studies demonstrated the tendency of abnormal tumorous gonads to form germline-derived teratoma [3]. Here, we demonstrate that cell differentiation decisions in the tumorous germline of animals deficient in the germline-specific translation repressor *gld-1*, can be regulated by neuronal cues that are sensitive to ER stress. Interestingly, we demonstrate that the propensity of the tumorous germline to differentiate into somatic cells rather than to maintain pluripotency is actively counteracted by neuronal cues of a newly-identified germline differentiation-inhibitory signaling circuit.

The germline differentiation-inhibitory neuronal circuit identified in this study includes the sensory neuron ASE, the interneuron AIY, and the motor neuron HSN, which synapses at the vulva. Furthermore, we identified the central signaling molecules required for the active suppression of GED by this neuronal circuit. While ASE communicates via the neuropeptide FLP-6, AIY communicates via the neurotransmitter acetylcholine and HSN communicates via serotonin. Deficiency in any of these neurons or signaling molecules comprising the germline differentiation-antagonizing circuit resulted in extreme ectopic differentiation of the germline in animals with a germline tumor **(see model, Fig. 6B).** Interestingly, additional reproduction-related decisions are determined non-autonomously by neuronal signals. For example, the ASI sensory neuron regulates germline apoptosis, germline mitosis and sperm activity [24, 30, 51]. Nevertheless, in tumorous *gld-1* animals, shifting germline fate from a pluripotent state to an ectopically differentiated somatic state is regulated by a distinct neuronal circuit, governed by the ASE sensory neuron. This implies that separate sets of neurons can provoke divergent responses in the *C. elegans* germline. ASE is a gustatory neuron, known mainly to mediate chemotaxis toward water-soluble cues, including salt ions such as Na^+^ and Cl^-^ [52]. While it may be puzzling why a salt-sensing neuron would control germline fate, previous studies demonstrated that the sensing of salt may be coupled to the presence of food [53]. In turn, food availability is an important determinant of reproductive physiology.

The most downstream signaling molecule identified in the neuronal circuit that controls germline pluripotency in the tumorous germline is the conserved neurotransmitter serotonin.

In *C. elegans*, serotonin causes dramatic behavioral effects including inhibition of locomotion, stimulation of egg laying and pharyngeal pumping and food-related behaviors [54–61]. While all these effects are modulated by the same molecule, they lead to different independent outcomes. Interestingly, none of the four dedicated *C. elegans* serotonin receptors is required for GED prevention. This may be due to redundancy or due to involvement of other, less specific, serotonin receptors in this circuit. Another layer of specificity may be encoded in the local levels and production site of serotonin. In support of this, serotonin production specifically by the HSN neuron is exclusively critical for preventing GED. Furthermore, migration-defective HSN does not effectively prevent GED, highlighting the importance for the correct location of the synapse of HSN near the vulva. Altogether, this suggests that the site and local concentrations of serotonin release play a key role in GED prevention, which differentiates this function from those of other serotonin signaling events. Furthermore, given the tumor-suppressive effects of GED [13] and consistent with studies which suggest that serotonin levels can play a crucial role in human cancer [62], this work supports a role for the serotoninergic system in tumor progression, rendering it a potential chemo-therapeutic target.

Similarly to GED induction by perturbations in any of the components of the germline differentiation-antagonising neuronal circuit, ER stress also induces GED within the tumorous gonads leading to suppression of the germline tumor [13]. Our present findings provide mechanistic insights to the underlying mechanism of ER stress-induced GED. Specifically, we show that ER stress-induced GED is not the consequence of ER stress *per se* within the germ cells, but rather the indirect result of activation of the IRE-1 endoribonuclease in the ASE neuron. We further show that the *flp-6* transcript has the structural requirements for RIDD, undergoes direct cleavage by IRE-1 *in-vitro*, and is destabilized by *ire-1 in-vivo*, indicating that the *flp-6* transcript is a target of RIDD. To the best of our knowledge, this is the first demonstration of a RIDD substrate in *C. elegans*. Since the *flp-6*transcript encodes the ASE-produced neuropeptide that controls the GED regulatory circuit, this disrupts the germline differentiation-antagonizing neuronal circuit, which can no longer prevent the acquisition of somatic fate by the germline tumor cells. This model is consistent with our previous findings that it is not the stress itself that regulates germ cell fate, but rather the activation of the ER stress sensor IRE-1, which in turn transduces an *xbp-1*-independent signal to promote GED and limit the progression of the germline tumor [13]. Altogether, this is the first evidence that neuronal circuits can be disrupted by RIDD to affect tumor progression by regulating crosstissue communication.

What could be the benefit in a germline differentiation-inhibitory neuronal circuit? We speculate that under physiological conditions, in a non tumorous background, such a circuit may be beneficial for preventing precocious differentiation of germline cells prior to fertilization. In the context of animals with a tumorous germline, we have previously demonstrated that acquisition of somatic fate by the tumorous germ cells limits the progression of the tumor by reducing its mitotic capacity and by restoring the ability of the cells to execute apoptosis [13]. The current identification of an ER stress-regulated neuronal circuit which controls this transition in a distal tumorous tissue suggests that targeting such neuronal circuits and targeting the RNase activity of IRE1 may be useful as therapeutic interventions for tumor suppression. These findings may be relevant to human tumors as well, since many human tumors, originating from a variety of major organs, are innervated, and this innervation is thought to contribute to the pathophysiology of cancer progression [63, 64].

## Acknowledgments

Some nematode strains were provided by the Caenorhabditis Genetics Center, which is funded by the NIH National Center for Research Resources and by Dr. Shohei Mitani, National Bioresource Project for the nematode, Tokyo Women’s Medical University School of Medicine, Japan. We thank Prof. Cori Bargmann (Rockefeller University, USA) for the HisCl related strains as well as for the Cre-Lox specific knockouts of *tph-1*. We thank Prof. Kang Shen (Stanford University, USA) for IRE-1 structure function strains (*wy782, wy762*). We thank Prof. Hannes E. Bulow (Albert Einstein College of Medicine, USA) for helpful discussions. We thank Prof. Dayong Wang (Southeast University, China) for *flp-6* rescue in ADF strain (*Psrh-142::flp-6; flp-6(ok3056)*). We thank Prof. Chris Li (The City College of New York, USA) for *flp-4(yn35)* mutant strain. We thank Dr. Jeniffer Israel-Cohen for the statistical analysis. This work was supported by: The Israel Science Foundation [grant number 1571/15 to S.H.K. The funders had no role in study design, data collection and analysis, decision to publish, or preparation of the manuscript.

## Author contributions

M.L.F, A.A and S.H.K. conceived and designed the experiments. M.L.F, R.S, A.L.T and M.S performed the experiments. M.L.F and Y.S generated plasmids and transgenic lines. A.A designed the *in-vitro* RIDD experiment and edited the manuscript. M.L.F and S.H.K. analyzed the data and wrote the manuscript.

## Materials and methods

### RNA interference

Bacteria expressing dsRNA were cultured overnight in LB containing tetracycline and ampicillin. Bacteria were plated on NGM plates containing 5mM IPTG and 50μg/ml carbenicillin. RNAi clone identity was verified by sequencing. Eggs were placed on plates and synchronized at day-0 (L4).The efficacy of the *tfg-1* RNAi was confirmed by the animals’ reduced body size [25] and by the induction of the ER stress reporter *Phsp-4::gfp* [24, 25]). The efficacy of the *ced-3* RNAi was confirmed by the lack of apoptotic corpses in the gonads. The efficacy of the *gld-1* RNAi was confirmed by the tumorigenicity of the gonads and by the absence of oocytes and embryos. For double or triple RNAi mixtures, the relative amount of each RNAi bacteria was kept equal between samples by growing the bacterial cultures overnight and then supplementing the relative amount of control RNAi as needed. Note that we have previously shown that GLD-1protein levels were efficiently and similarly reduced in all RNAi combinations involving *gld-1* RNAi including single RNAi treatment, double RNAi treatment and triple RNAi treatment (See Figure 1 – figure supplement 2 in [13]).

### Fluorescence microscopy and quantification

To follow expression of fluorescent transgenic markers, transgenic animals were anaesthetized on 2% agarose pads containing 2 mM levamisol. Images were taken with a CCD digital camera using a Nikon 90i fluorescence microscope. For each trial, exposure time was calibrated to minimize the number of saturated pixels and was kept constant through the experiment. The NIS element software was used to quantify mean fluorescence intensity as measured by intensity of each pixel in the selected area within the gonad.

To determine the fraction of the gonad area occupied by ectopic cells, day-4 animals were fixed and stained with DAPI. The NIS element software was used to manually select and quantify the gonad area as well as the area within the gonad that was occupied by abnormal DAPI-stained nuclei in the animals. Percent of gonad area occupied by large nuclei is the ratio of these two paired measurements.

### Statistical analysis

Error bars represent the standard error of the mean (SEM) of independent biological replicates unless indicated otherwise. For qRT-PCR, P-values were calculated using the unpaired Student’s t test. For a simple comparison between two data sets, P values were determined using unpaired Student’s T-test, assuming unequal variances. For multiple comparisons, between multiple data sets, groups and/or treatments were compared using one or two-way ANOVA followed by a Tukey’s post hoc analysis. Normality of residuals assumption was assessed through with residuals plots. See table S2 for statistical data. Significance threshold was set as P<0.01 and marked with an asterisk. Significant changes beyond 2-fold change is marked by three asterisks.

### Quantitative RT-PCR analysis for *flp-6*

Animals were raised at 20°C until day-1 of adulthood. On day-1, animals were collected for RNA extraction. RNA extraction, purification and reverse transcription were carried out using standard protocols. Real-time PCR was performed using Maxima SYBR (Fermentas) in a StepOnePlus instrument. Transcript levels of *act-1* were used for normalization.

***act-1:*** 5-CCAATCCAAGAGAGGTATCCTTAC-3’ and

5’-CATTGTAGAAGGTGTGATGCCAG-3’

***flp-6:*** 5’ - GTGAAGTGGAGAGAGAAATGATGA - 3’ and

5-CCGCTACTTCTCTTTCCAAAACG-3’

### DAPI staining

Adult worms (at the ages of either day1, day 4, day 5 or day 10) were collected from agar plates and were washed with M9 solution to remove *E. coli* bacteria. Permeabilization of the animals was performed by freezing them at −80°C. For fixation, worms were washed with cold methanol and incubated for 15 minutes at −20°C. The fixed animals were then washed twice with PBSTx1 and stained with 1μg/mL DAPI (4’,6-diamidino-2-phenylindole)(Sigma) solution for 30 minutes. Worms were washed two times with PBSTx1 to remove excess staining and observed under the fluorescent microscope to quantify ectopic differentiated nuclei within the animals’ gonads.

### Serotonin treatment

Nematodes were grown and assayed at room temperature on standard NGM seeded with *E. coli* strain OP50 as a food source till day-1 of adulthood. For drug experiments, 5-hydroxytryptamine creatinine sulfate complex (Sigma) was added to NGM agar plates to a final concentration of 20 mM. Day-1 animals were grown till day-4 on the 20 mM 5-HT plates and washed for DAPI staining procedure at day-4 of adulthood.

### Histamine treatment

As previously described [46], eggs from four histamine-inducible interneurons silencing mutants were grown on non-histamine OP50 seeded plates until L4-stage. At L4 stage, worms were moved to histamine containing plates (represented by black bars in Fig. 4A as histamine +) or to non-histamine plates (represented by gray bars in Fig. 4A as histamine-) until Day 4 of adulthood. At day-4 of adulthood worms were examined for ectopic large nuclei within the gonads using DAPI staining. 1 M histamine (HA) stock was prepared by Dissolving 1.85 g histamine dihydrochloride (Sigma-Aldrich H7250) per 10 mL water. Experiments were performed in Plates contacting a final concentration of 10mM histamine.

### RNA cleavage assay

1 μg of T7 RNA generated was digested at room temperature by 1 μg of human IRE-1 α KR recombinant protein (~0.8 μM) for 15 min in RNA cleavage buffer [HEPES pH7.5 20 mM; potassium acetate 50 mM; magnesium acetate 1 mM; TritonX-100 0.05% (v/v)]. The total volume of the reaction is 25 μl. The digestion was then complemented by an equal volume of formamide and heated up at 70°C for 10 min to linearize the RNA. After linearization, the mixture was immediately placed on ice for 5 min, and then 20 μl was run on 3% agarose gel at 160 V for 50 min. If inhibitor was used, it was incubated with IRE-1α KR for 40 min on ice prior to RNA digestion. Gels were visualized on either a BioRad Molecular Imager ChemiDoc ZRS+. The T7 RNA transcripts was generated from cDNA templates of *flp-6*. cDNA was amplified, adding T7 sequence to forward (T7-FLP6-f) and BGH sequence to reverse (BGH-FLP6-r) primers, and subsequently in-vitro transcribed using HiScribe™ T7 Quick High Yield RNA Synthesis Kit from NEB (#E2050S). The IRE-1 RNase-specific inhibitor 4μ8c was used at 5 μM.

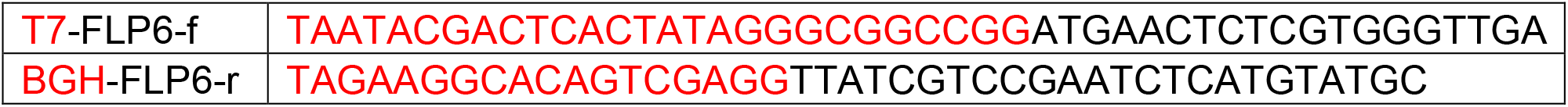

### Strains

The following lines were used in this study: N2, SHK124: *rrf-1(pk1417) I*, CF2473: *ire-1(ok799) II*, SHK27: *ire-1(ok799) II; biuEx2[pRF4(rol-6(su1006)); Pire-1::ire-1(cDNA)], SHK185: ire-1(ok799) II; biuEx49[pRF4(rol-6(su1006)): Prgef-1::ire-1(cDNA)]*, SHK14: *ire-1(ok799) II; biuEx5[pRF4(rol-6(su1006)); Pmyo-3::ire-1(cDNA)]*, YES85*: ire-1(ok799) II; biuEx50[pRF4(rol-6(su1006)); Pges-1::ire-1(cDNA)]*, SHK4*: biuEx2[pRF4(rol-6(su1006)); Pire-1::ire-1(cDNA)]*, SHK182: *biuEx49[pRF4(rol-6(su1006)); Prgef-1::ire-1(cDNA)]*, SHK8: *biuEx5[pRF4(rol-6(su1006)); Pmyo-3::ire-1(cDNA)]*, SHK282: *biuEx50[pRF4(rol-6(su1006)); Pges-1::ire-1(cDNA)]*, SHK268: *biuEx11[Pgrd-10::gfp; Pche-12::ire-1(cDNA)]; ire-1(ok799) II*, SHK271*: biuEx14[Pgrd-10::gfp; Peat-4::ire-1(cDNA)]; ire-1(ok799) II*, SHK270*: biuEx13[Pgrd-10::gfp; Pdat-1::ire-1(cDNA)]; ire-1(ok799) II*, SHK266*: biuEx10[Pgrd-10::gfp; Punc-25::ire-1(cDNA)]; ire-1(ok799) II*, SHK15: *biuEx4[pRF4(rol-6(su1006)); Pdaf-28::ire-1(cDNA)]; ire-1(ok799) II*, SHK256*: biuEx51[pRF4(rol-6(su1006)); Pdaf-7::ire-1]; ire-1(ok799) II*, SHK592*: biuEx57[pRF4(rol-6(su1006)); Pche-1::ire-1]; ire-1(ok799) II*, SHK593: *biuEx58[pRF4(rol-6(su1006)); Pops-1::ire-1]; ire-1(ok799) II, OH10434: Pche-1::mCherry::che-1 3UTR; pRF4(rol-6(su1006)). DA572: eat-4(ad572), MT6308: eat-4(ky5), NY119: flp-4(yn35); VC2324: flp-6(ok3056)V, FX02427; flp-13(tm2427), PT505: flp-20(pk1596) X, VC1982: flp-25(gk1016), FX03023: nlp-3(tm3023), RB1609: nlp-5(ok1981) II, FX1880: nlp-14(tm1880), FX1888: ins-1(tm1888), RB2594: ins-22(ok3616), FX14756: ins-26(tm1983), FX06109: ins-32(tm6109), RB982: flp-21(ok889) V, RB1340: nlp-1(ok1469), FX02984: nlp-7(tm2984), VC1309: nlp-8(ok1799) I, nlp-10(tm6232), VC1063: nlp-15(ok1512) I, SHK497: biuEx52[pRF4(rol-6(su1006)); Pche-1::flp-6(cDNA)]; flp-6(ok3056), SHK498: biuEx53[pRF4(rol-6(su1006)); Psrh-142::flp-6]; flp-6(ok3056), PR672: che-1(p672), PR674: che-1(p674), MT633: lin-11(n389) I; him-5(e1467)V, MT1196: lin-11(n566) I, CF2260: Zcls4[Phsp-4::gfp] V, SHK314: Zcls4[Phsp-4::gfp] V; flp-6(ok3056), SHK315: ire-1(ok799) II, flp-6(ok3056), NY2067: ynIs67[Pflp-6::gfp] III; him-5(e1490) V, SHK403: ynIs67[Pflp-6::gfp] III; ire-1(ok799) II, SHK474: biuEx54[pRF4(rol-6(su1006)); Pche-1::flp-6(cDNA)::gfp], SHK491: biuEx52[pRF4(rol-6(su1006)); Pche-1::flp-6(cDNA)]; ire-1(ok799) II, TV13656: ire-1(wy782), TV13763: ire-1(wy762), NY7: flp-1(yn2) IV, NY16: flp-1(yn4) IV, AX1410: flp-18(db99) X, VC2016: flp-18(gk3063) X, PR1152: cha-1(p1152) IV, SHK605: biuEx55[pRF4(rol-6(su1006)); Pttx-3::cha-1]; cha-1(p1152) IV, GG201: ace-2(g72); ace-1(p1000), PR1300: ace-3(dc2), CB113: unc-17(e113), SHK586: biuEx56[pRF4(rol-6(su1006)); Pttx-3::unc-17]; unc-17(e113), MT9668: mod-1(ok103) V, DA1814:ser-1(ok345) X, AQ866: ser-4 (ok512) III, RB1585: ser-7(ok1944) X, MT1446: her-1(n695) V, MT5825: sem-4(n1971) I, MT3969: mig-1(n1652) I, NF69:mig-19(k142) II, NF78: mig-20(k148) X, SHK492: flp-6(ok3056); sem-4(n1971) I, SHK490: flp-6(ok3056); her-1(n695) V, FX30280: flp-5(tm10075), LC33: bas-1(tm351) III CX13228: tph-1(mg280) II; kySi56[tph-1 genomic kyEx4107[egl-rescue] IV, CX13576: tph-1(mg280) II; kySi56[ tph-1 genomic rescue] IV 6::nCre], CX13571: tph-1(mg280) II; kySi56[ tph-1 genomic rescue] IV; kyEx4077[srh-142::nCre], CX13574: tph-1(mg280) II; kySi56[ tph-1 genomic rescue] IV; kyEx4081[ops-1::nCre], CX14909: kyEx4925 [ttx-3::hisCl1*::sl2::GFP; myo-3::mCherry], CX14849: kyEx4867 [ins-1::HisCl1::sl2mCherry; unc-122::GFP]*, CX14908: *kyEx4924 [inx-1::hisCl1*::sl2::GFP; myo-3::mCherry]*, CX16069: kyEx5493 [pNP443 *(glr-3::HisCl1::SL2::mCherry); elt-2:mCherry]*. DP132: *edIs6 (punc-119::GFP) IV, SHK659:che-1(p679) I; edIs6 (punc-119::GFP) IV*, HK361: *edIs6 (punc-119::GFP) IV; flp-6(ok3056)*, SHK660: *cha-1(p1152) edIs6 (punc-119::GFP) IV*, SHK661: *sem-4(n1378) I; edIs6 (punc-119::GFP) IV*, SHK662: *tph-1(mg280) II; edIs6 (punc-119::GFP) IV, SHK663: Zcls4[Phsp-4::gfp] V; otIs232 [che-1p::mCherry(C. elegans-optimized)::che-1 3’UTR + rol-6(su1006)]*, OH10434 *otIs232 [che-1p::mCherry(C. elegans-optimized)::che-1 3’UTR + rol-6(su1006)]*

### Molecular cloning

*Pire-1::ire-1, Prgef-1::ire-1, Pdaf-7::ire-1, Pdaf-28::ire-1* and *Pmyo-3::ire-1* plasmids have been previously described [24]. *Psrh-142::flp-6* has been previously described [65].

*Peat-4::ire-1, Punc-25::ire-1, Pche-12::ire-1* and *Pdat-1::ire-1* plasmids have been genrated as follows: the *ire-1* coding sequence was amplified from cDNA and cloned into the *Nhe*I and *Kpn*I sites in the L3691 plasmid. Genomic DNA-amplified promoters were inserted upstream of the *ire-1* coding sequence (CDS) as follows: *Peat-4* (~3 kb), *Punc-25* (~1.6 kb), *Pche-12* (~ 1 kb) and *Pdat-1* (~0.7 kb) were inserted in the SphI/NotI restriction sites. *Pges-1* (~3 kb) was inserted in the SphI/KpnI restriction sites.

To generate *Pche-1::flp-6* the *flp-6* coding sequence was amplified from cDNA and cloned into the XmaI and KpnI sites in the Andy-Pire L3691 vector. *Pche-1* (ASE specific, ~1.4 kb) promoter was cloned into NotI and XmaI sites upstream to *flp-6* cDNA, replacing the original *mec-7* promoter.

To generate *Pttx-3::cha-1* and *Pttx-3::unc-17* the *cha-1* coding sequence and the *unc-17* coding sequence were amplified from cDNA and cloned into in the *KpnI/SphI* sites downstream of the *Pttx-3*^promB^ regulatory element in the *Pttx-3::ire-1* vector, previously described [24], replacing the *ire-1* coding sequence.

Germline transformations were performed by injection of 25 ng/ml plasmids and 100 ng/ml of *rol-6(su1006)* as a co-transformation marker.

---

**Figure 2 – figure supplement 1.**
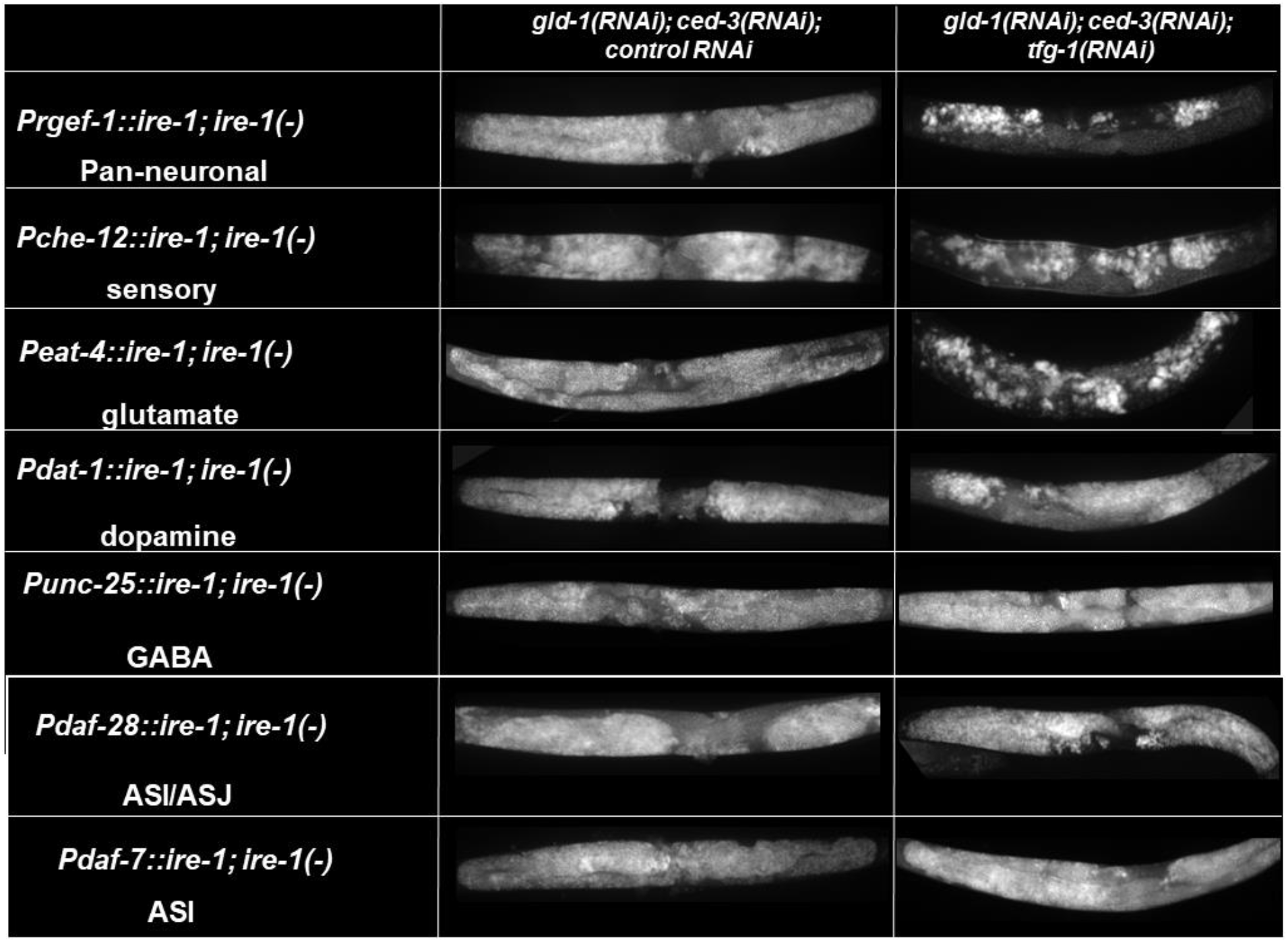
ER stress-induced GED is controlled by sensory neuronal IRE-1. Representative micrographs of whole body (x100) DAPI-stained day-4 *ire-1(-)* transgenic animals, each expressing an *ire-1*-rescuing transgene driven by the indicated promoters, treated with either a mixture of control, *gld-1* and *ced-3* RNAi or with a mixture of *tfg-1, gld-1* and *ced-3* RNAi. (n= 210 gonads per genotype, N=4). See Fig. 2A for quantification.

**Figure 3 – figure supplement 1.**
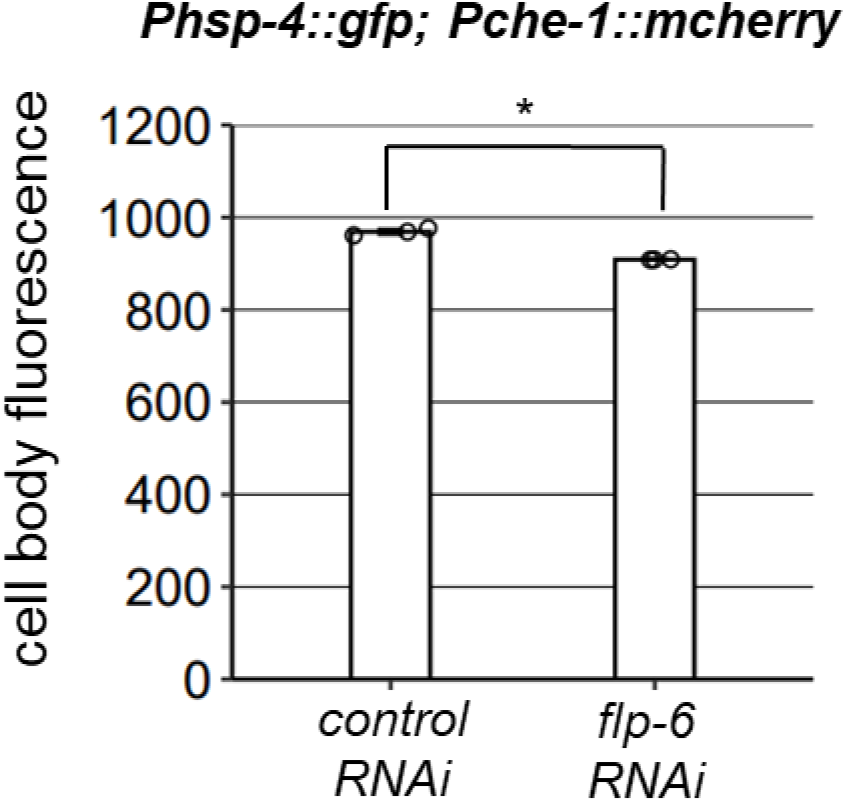
*Phsp-4* activity in the ASE neurons is not altered by *flp-6* RNAi. The fluorescence levels of the *Phsp-4::GFP* ER stress reporter were not increased by *flp-6* RNAi treatment (Student’s T-test P=0.0056, n=95 animals per genotype, N=3). The selected areas for fluorescence measurements were manually selected based on their overlap with the *Pche-1::mcherry* marker.

**Figure 3 – figure supplement 2.**
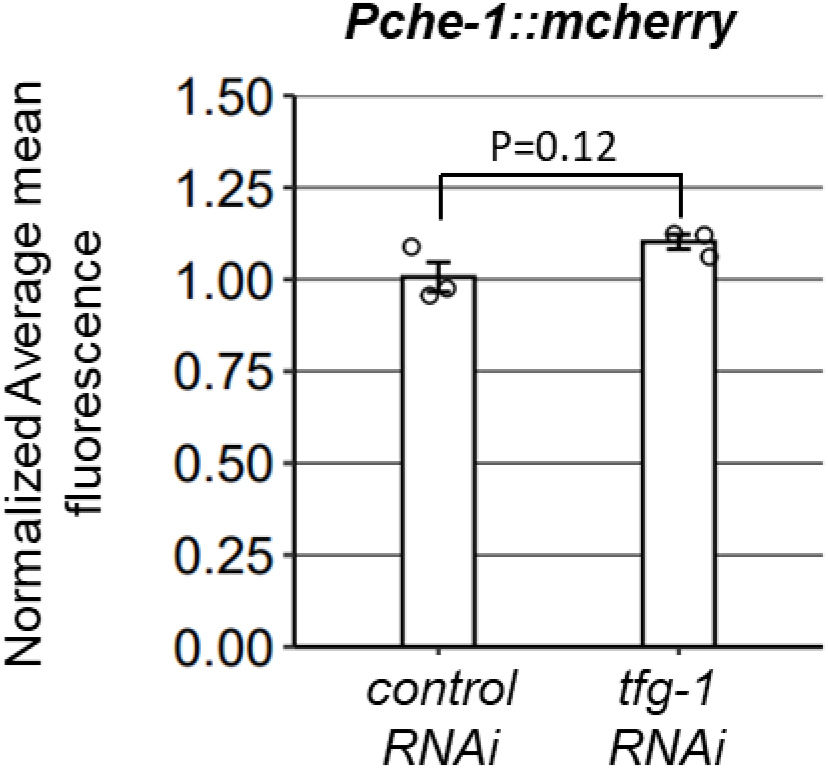
*Pche-1* activity is not altered by ER stress. The fluorescence levels of the *Pche-1::mcherry* transcriptional reporter were not significantly altered by *tfg-1* RNAi treatment (Student’s T-test P=0.1258, n=120 animals per genotype, N=3).

**Figure 3 – figure supplement 3.**
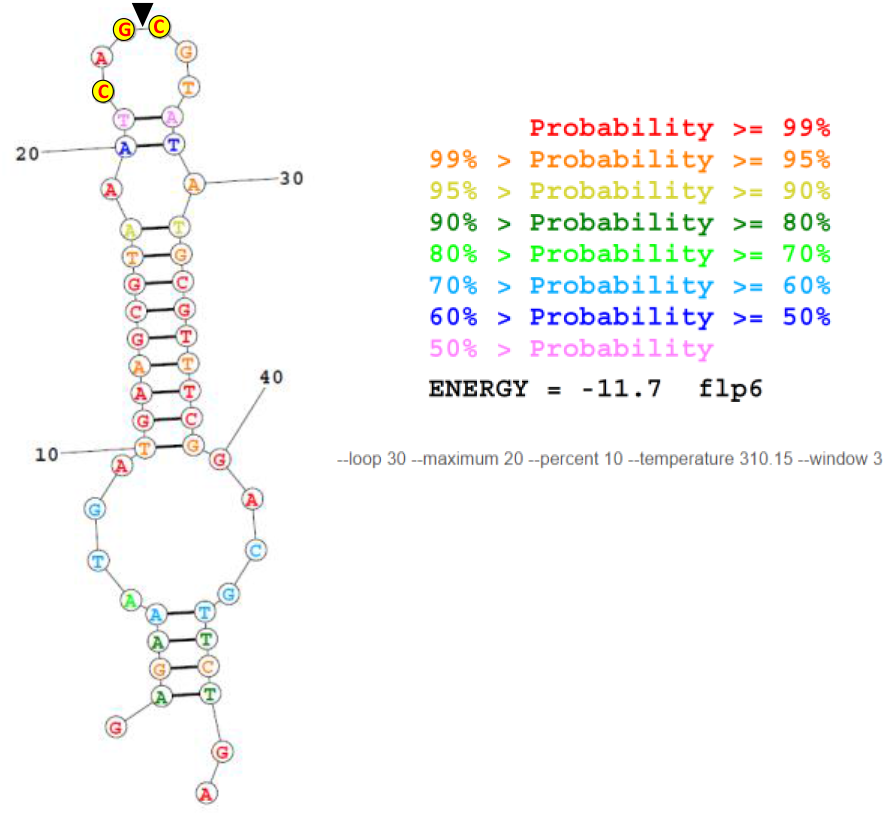
The lowest free energy predicted secondary RNA structure for the given fragment of the *flp-6* transcript sequence as calculated by the RNA structure web server [66]. The conserved residues of in the nucleotide loop of putative RIDD substrates [39] are highlighted in yellow. Black triangle marks putative cleavage site by IRE1.

**Figure 5 – figure supplement 1.**
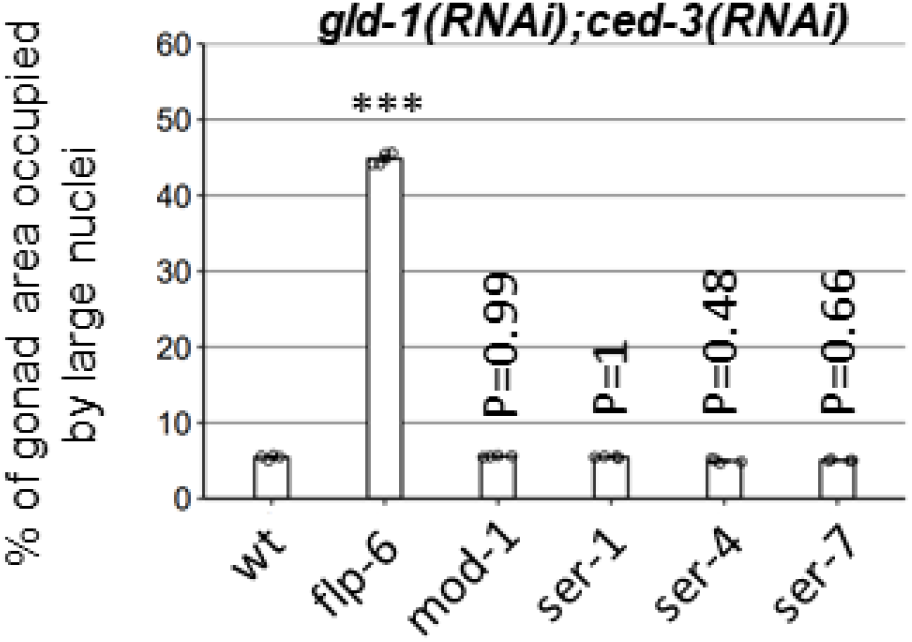
Four major serotonin receptor mutants are not required for GED in *gld-1;ced-3* deficient animals. Bar graphs present percentage of gonad area occupied by DAPI-labeled ectopic cells in day-4 animals treated with *gld-1* and *ced-3* RNAi (n= 140 gonads per genotype, N=4). Asterisks mark one-way ANOVA followed by Tukey’s post hoc analysis of p < 0.001 relative to wild-type animals treated with *gld-1* and *ced-3* RNAi. *mod-1,ser-1, ser-4 and ser-7* deficiencies were obtained by *ok103, ok345, ok512, ok1944* mutations respectively. Triple asterisks mark significant results resulting in a 2-fold change or more.

**Table S1 –.** List of sensory neurons included under the che-1 promoter

| List of sensory neurons included under the *che-12* promoter |  |  |
| --- | --- | --- |
| Neuron Name | Neurotransmitter | Neuropeptides |
| ASE | Glutamate | flp-4, flp-6, flp-13, flp-20, flp-25, ins-1, ins-22, ins-26, ins-32, nlp-3, nlp-5, nlp-14 |
| AWC | Glutamate | nlp-1 |
| ASH | Glutamate | flp-21, ins-1, nlp-3, nlp-15 |
| ASK | Glutamate | flp-21, nlp-8, nlp-10, nlp-14 |
| ASG | Glutamate | flp-6, flp-13, flp-22, ins-1 |
| ADL | Glutamate | flp-4, flp-21, nlp-7, nlp-8, nlp-10 |
| AQR | Glutamate | - |
| PQR | Glutamate | - |
| PHA | Glutamate | nlp-7, nlp-14 |
| PHB | Glutamate | nlp-1 |
| AWA | - | - |
| AWB | Acetylcholine | nlp-3, nlp-9 |
| ASI | - | ,daf-28, flp-2, flp-10, flp-21, ins-1, ins-3, ins-4, ins-6, ins-7, ins-9, ins-22, ins-26, ins-32, nlp-1, nlp-5, nlp-6, nlp-7, nlp-9, nlp-14, nlp-18, nlp-24, nlp-27, pdf-1 |
| ADF | Acetylcholine, Serotonin | flp-6, ins-1, nlp-3 |
| ASJ | - | nlp-3 |
| URX | Acetylcholine | flp-8, flp-10, flp-11, flp-19 |

**Table S2 –.** Statistical data

|  |  |  |  |  |  |
| --- | --- | --- | --- | --- | --- |
| **Fig.1A : One-way ANOVA followed by Tukey's post hoc analysis** |  |  |  |  |  |
| all animals treated with gld-1 and ced-3 RNAi |  |  | degrees of freedom | p-value | F |
|  |  |  | 3(20) | <0.0001 | 29740.9165 |
| comparison | p-value | difference |  |  |  |
| tfg-1(RNAi) vs. control(RNAi) | P<0.001 | 22.2963 |  |  |  |
| rff-1(-); tfg-1(RNAi) vs. rff-1(-); control(RNAi) | P<0.001 | 1.1757 |  |  |  |
| **Fig.1B : One-way ANOVA followed by Tukey's post hoc analysis** |  |  |  |  |  |
| g.c.t indictaes treatment with gld-1; ced-3 and tfg-1 RNAi. |  |  | degrees of freedom | p-value | F |
| g.c.c indictaes treatment with gld-1; ced-3 and control RNAi |  |  | 6(21) | <0.0001 | 65652.2081 |
| comparison | p-value | difference |  |  |  |
| tfg-1(RNAi) g.c.t vs. control(RNAi) g.c.c | P<0.001 | 19.001 |  |  |  |
| ire-1(-); Pire-1::ire-1 g.c.t vs. ire-1(-) g.c.t | P<0.001 | 19.4796 |  |  |  |
| ire-1(-); Prgef-1::ire-1 g.c.t vs. ire-1(-) g.c.t | P<0.001 | 23.9432 |  |  |  |
| ire-1(-); Pmyo-3::ire-1 g.c.t vs. ire-1(-) g.c.t | P<0.001 | -0.4249 |  |  |  |
| ire-1(-); Pges-1::ire-1 g.c.t vs. ire-1(-) g.c.t | P<0.001 | -0.4624 |  |  |  |
| tfg-1(RNAi) g.c.t vs. ire-1(-); Pire-1::ire-1 g.c.t | 0.0437 | -0.2078 |  |  |  |
| **Fig.1C : One-way ANOVA followed by Tukey's post hoc analysis** |  |  |  |  |  |
| all animals treated with gld-1; ced-3 RNAi |  |  | degrees of freedom | p-value | F |
|  |  |  | 3(12) | <0.0001 | 35987.922 |
| comparison | p-value | difference |  |  |  |
| Pmyo-3::ire-1 vs. Pges-1::ire-1 | 0.0302 | -0.3205 |  |  |  |
| Prgef-1::ire-1 vs. Pges-1::ire-1 | P<0.001 | 23.2125 |  |  |  |
| Pire-1::ire-1 vs. Pges-1::ire-1 | P<0.001 | 22.1319 |  |  |  |
| Pire-1::ire-1 vs. Pmyo-3::ire-1 | P<0.001 | 22.4523 |  |  |  |
| Pire-1::ire-1 vs. Prgef-1::ire-1 | P<0.001 | -1.0807 |  |  |  |
| **Fig.2A : two-way ANOVA followed by Tukey's post hoc analysis** |  |  |  |  |  |
| g.c.t indictaes treatment with gld-1; ced-3 and tfg-1 RNAi. |  |  | degrees of freedom | p-value | F |
| g.c.c indictaes treatment with gld-1; ced-3 and control RNAi |  |  | 6(42) | <0.0001 | 3052.5167 |
| comparison | p-value | difference |  |  |  |
| Pche-12::ire-1; ire-1(-):g.c.t vs. Pche-12::ire-1; ire-1(-):g.c.c | P<0.001 | 24.8178 |  |  |  |
| Pdaf-28::ire-1; ire-1(-):g.c.t vs. Pdaf-28::ire-1; ire-1(-):g.c.c | 0.9986 | 0.2463 |  |  |  |
| Pdaf-7::ire-1; ire-1(-):g.c.t vs. Pdaf-7::ire-1; ire-1(-):g.c.c | 1 | 0.0944 |  |  |  |
| Pdat-1::ire-1; ire-1(-):g.c.t vs. Pdat-1::ire-1; ire-1(-):g.c.c | 1 | 0.1778 |  |  |  |
| Peat-4::ire-1; ire-1(-):g.c.t vs. Peat-4::ire-1; ire-1(-):g.c.c | P<0.001 | 24.0679 |  |  |  |
| Prgef-1::ire-1; ire-1(-):g.c.t vs. Prgef-1::ire-1; ire-1(-):g.c.c | P<0.001 | 25.7532 |  |  |  |
| Punc-52::ire-1; ire-1(-):g.c.t vs. Punc-52::ire-1; ire-1(-):g.c.c | 0.9983 | 0.2516 |  |  |  |
| **Fig.2B : two-way ANOVA followed by Tukey's post hoc analysis** |  |  |  |  |  |
| g.c.t indictaes treatment with gld-1; ced-3 and tfg-1 RNAi. g.c.c indictaes treatment with gld-1; ced-3 and control RNAi |  |  | degrees of freedom | p-value | F |
|  |  |  | 1(8) | 0.0061 | 13.6368 |
| comparison | p-value | difference |  |  |  |
| eat-4(ad572):g.c.t vs. eat-4(ad572):g.c.c | P<0.001 | 25.1305 |  |  |  |
| eat-4(ky5):g.c.t vs. eat-4(ky5):g.c.c | P<0.001 | 26.5342 |  |  |  |
| **Fig.2C : two-way ANOVA followed by Tukey's post hoc analysis** |  |  |  |  |  |
| g.c.t indicat treatment with gld-1; ced-3 and tfg-1 RNAi. |  |  |  |  |  |
| g.c.c indicat treatment with gld-1; ced-3 and control RNAi |  |  |  |  |  |
|  |  |  | **degrees of freedom** | **p-value** | **F** |
|  |  |  | 19(120) | <0.0001 | 2731.0279 |
| **comparison** | **p-value** | **difference** |  |  |  |
| **wt:g.c.c-wt:g.c.t** | **P<0.001** | **41.3852** |  |  |  |
| wt:g.c.c vs. flp-4(-):g.c.c | P<0.001 | 1.0723 |  |  |  |
| **wt:g.c.c vs. flp-6(-):g.c.c** | **P<0.001** | **43.1677** |  |  |  |
| wt:g.c.c vs. flp-13(-):g.c.c | P<0.001 | 2.9964 |  |  |  |
| wt:g.c.c vs. flp-20(-):g.c.c | 0.7275 | 0.4275 |  |  |  |
| wt:g.c.c vs. flp-21(-):g.c.c | P<0.001 | 3.3928 |  |  |  |
| wt:g.c.c vs. flp-22(-):g.c.c | P<0.001 | 0.9706 |  |  |  |
| wt:g.c.c vs. flp-25(-):g.c.c | 0.8395 | 0.4006 |  |  |  |
| wt:g.c.c vs. nlp-1(-):g.c.c | P<0.001 | 1.569 |  |  |  |
| wt:g.c.c vs. nlp-3(-):g.c.c | P<0.001 | 1.9358 |  |  |  |
| wt:g.c.c vs. nlp-5(-):g.c.c | P<0.001 | 1.8992 |  |  |  |
| wt:g.c.c vs. nlp-7(-):g.c.c | P<0.001 | 1.0701 |  |  |  |
| wt:g.c.c vs. nlp-8(-):g.c.c | P<0.001 | 2.9725 |  |  |  |
| wt:g.c.c vs. nlp-10(-):g.c.c | P<0.001 | 2.9235 |  |  |  |
| wt:g.c.c vs. nlp-14(-):g.c.c | P<0.001 | 2.957 |  |  |  |
| wt:g.c.c vs. nlp-15(-):g.c.c | P<0.001 | 3.1653 |  |  |  |
| wt:g.c.c vs. ins-1(-):g.c.c | P<0.001 | 1.1331 |  |  |  |
| wt:g.c.c vs. ins-22(-):g.c.c | 1 | 0.0382 |  |  |  |
| wt:g.c.c vs. ins-26(-):g.c.c | P<0.001 | 3.4378 |  |  |  |
| wt:g.c.c vs. ins-32(-):g.c.c | P<0.001 | 1.8764 |  |  |  |
| wt:g.c.t vs. flp-4(-):g.c.t | P<0.001 | 1.8847 |  |  |  |
| wt:g.c.t vs. flp-6(-):g.c.t | P<0.001 | 3.5812 |  |  |  |
| wt:g.c.t vs. flp-13(-):g.c.t | P<0.001 | 2.4021 |  |  |  |
| wt:g.c.t vs. flp-20(-):g.c.t | P<0.001 | 3.3803 |  |  |  |
| wt:g.c.t vs. flp-21(-):g.c.t | P<0.001 | 2.4342 |  |  |  |
| wt:g.c.t vs. flp-22(-):g.c.t | P<0.001 | 2.9644 |  |  |  |
| wt:g.c.t vs. flp-25(-):g.c.t | P<0.001 | 2.4219 |  |  |  |
| wt:g.c.t vs. nlp-1(-):g.c.t | P<0.001 | 2.9715 |  |  |  |
| wt:g.c.t vs. nlp-3(-):g.c.t | P<0.001 | 3.4255 |  |  |  |
| wt:g.c.t vs. nlp-5(-):g.c.t | P<0.001 | 1.4221 |  |  |  |
| wt:g.c.t vs. nlp-7(-):g.c.t | P<0.001 | 5.4199 |  |  |  |
| wt:g.c.t vs. nlp-8(-):g.c.t | P<0.001 | 4.3791 |  |  |  |
| wt:g.c.t vs. nlp-10(-):g.c.t | P<0.001 | 2.4409 |  |  |  |
| wt:g.c.t vs. nlp-14(-):g.c.t | P<0.001 | 2.4292 |  |  |  |
| wt:g.c.t vs. nlp-15(-):g.c.t | P<0.001 | 4.3582 |  |  |  |
| wt:g.c.t vs. ins-1(-):g.c.t | P<0.001 | 4.4275 |  |  |  |
| wt:g.c.t vs. ins-22(-):g.c.t | P<0.001 | 5.3754 |  |  |  |
| wt:g.c.t vs. ins-26(-):g.c.t | P<0.001 | 4.4472 |  |  |  |
| wt:g.c.t vs. ins-32(-):g.c.t | P<0.001 | 1.4919 |  |  |  |
| **Fig.2D : Student's T-test** |  |  |  |  |  |
| all animals treated with gld-1; ced-3 RNAi |  |  |  |  |  |
| **comparison** | **p-value** | **difference** |  |  |  |
| **control(RNAi) vs. flp-6(RNAi)** | **P<0.001** | **39.0326** |  |  |  |
| **Fig.2E : One-way ANOVA followed by Tukey's post hoc analysis** |  |  |  |  |  |
| all animals treated with gld-1; ced-3 RNAi |  |  |  |  |  |
|  |  |  | **degrees of freedom** | **p-value** | **F** |
|  |  |  | 7(24) | <0.0001 | 11055.9311 |
| **comparison** | **p-value** | **difference** |  |  |  |
| **wt vs. flp-6(-)** | **P<0.001** | **40.0835** |  |  |  |
| **wt vs. che-1(p672)** | **P<0.001** | **35.9091** |  |  |  |
| **wt vs. che-1(p674)** | **P<0.001** | **36.8292** |  |  |  |
| wt vs. lin-11(n389) | 1 | 0.0242 |  |  |  |
| wt vs. lin-11(n566) | 0.0463 | 0.8995 |  |  |  |
| **Pche-1::flp-6; flp-6(-) vs. flp-6(-)** | **P<0.001** | **40.0764** |  |  |  |
| Psrh-142::flp-6; flp-6(-) vs. flp-6(-) | P<0.001 | 2.7991 |  |  |  |
| wt vs. Pche-1::flp-6; flp-6(-) | 1 | 0.007 |  |  |  |
| **wt vs. Psrh-142::flp-6; flp-6(-)** | **P<0.001** | **37.2843** |  |  |  |
| **Fig.3A: Student's T-test** |  |  |  |  |  |
| Phsp-4::gfp transgenic animals |  |  |  |  |  |
| **comparison** | **p-value** | **difference** |  |  |  |
| Phsp-4::gfp vs. Phsp-4::gfp; flp-6(-) | 0.4948 | 0.0982 |  |  |  |
| **Fig.3B : One-way ANOVA followed by Tukey's post hoc analysis** |  |  |  |  |  |
| all animals treated with gld-1; ced-3 RNAi |  |  |  |  |  |
|  |  |  | **degrees of freedom** | **p-value** | **F** |
|  |  |  | 3(12) | <0.0001 | 29409.8165 |
| **comparison** | **p-value** | **difference** |  |  |  |
| wt vs. ire-1(-) | 0.0203 | 0.6115 |  |  |  |
| **wt vs. flp-6(-)** | **P<0.001** | **37.1224** |  |  |  |
| **ire-1(-); flp-6(-) vs. ire-1(-)** | **P<0.001** | **36.5806** |  |  |  |
| ire-1(-); flp-6(-) vs. flp-6(-) | P<0.001 | 1.1533 |  |  |  |
| **Fig.3E : One-way ANOVA followed by Tukey's post hoc analysis** |  |  |  |  |  |
| Pflp-6::gfp transgenic animals |  |  |  |  |  |
|  |  |  | **degrees of freedom** | **p-value** | **F** |
|  |  |  | 3(12) | 0.0306 | 4.1761 |
| **comparison** | **p-value** | **difference** |  |  |  |
| tfg-1(RNAi) vs. control(RNAi) | 0.7487 | 0.017 |  |  |  |
| ire-1(-); tfg-1 RNAi vs. ire-1(-); control RNAi | 0.6933 | 0.0187 |  |  |  |
| ire-1(-); control RNAi-control(RNAi) | 0.1731 | 0.0376 |  |  |  |
| tfg-1(RNAi)-ire-1(-); tfg-1 RNAi | 0.1483 | 0.0392 |  |  |  |
| **Fig.3F : One-way ANOVA followed by Tukey's post hoc analysis** |  |  |  |  |  |
| Pche-1::flp-6::gfp transgenic animals |  |  |  |  |  |
|  |  |  | **degrees of freedom** | **p-value** | **F** |
|  |  |  | 3(12) | <0.0001 | 3354.3839 |
| **comparison** | **p-value** | **difference** |  |  |  |
| tfg-1(RNAi) vs. control(RNAi) | P<0.001 | 2.7253 |  |  |  |
| ire-1(-); tfg-1 RNAi vs. ire-1(-); control RNAi | P<0.001 | 0.8648 |  |  |  |
| **Fig.3G : two-way ANOVA followed by Tukey's post hoc analysis** |  |  |  |  |  |
| g.c.t indicat treatment with gld-1; ced-3 and tfg-1 RNAi.<br>g.c.c indicat treatment with gld-1; ced-3 and control RNAi |  |  |  |  |  |
|  |  |  | **degrees of freedom** | **p-value** | **F** |
|  |  |  | 3(32) | <0.0001 | 17452.9053 |
| **comparison** | **p-value** | **difference** |  |  |  |
| **wt:g.c.t- vs. wt:g.c.c** | **P<0.001** | **32.0115** |  |  |  |
| ire-1(ok799):g.c.t vs. ire-1(ok799):g.c.c | 0.0033 | 0.5236 |  |  |  |
| ire-1(wy782):g.c.t vs. ire-1(wy782):g.c.c | 0.0612 | 0.3835 |  |  |  |
| ire-1(wy762):g.c.t vs. ire-1(wy762):g.c.c | 0.3963 | 0.2639 |  |  |  |
| **Fig.3H : two-way ANOVA followed by Tukey's post hoc analysis** |  |  |  |  |  |
| g.c.t indicat treatment with gld-1; ced-3 and tfg-1 RNAi.<br>g.c.c indicat treatment with gld-1; ced-3 and control RNAi |  |  |  |  |  |
|  |  |  | **degrees of freedom** | **p-value** | **F** |
|  |  |  | 3(16) | <0.0001 | 14118.0507 |
| **comparison** | **p-value** | **difference** |  |  |  |
| **wt:g.c.t vs. wt:g.c.c** | **P<0.001** | **26.4023** |  |  |  |
| ire-1(-):g.c.t vs. ire-1(-):g.c.c | 0.6641 | 0.2418 |  |  |  |
| Pops-1::ire-1; ire-1(-):g.c.t vs. Pops-1::ire-1; ire-1(-):g.c.c | 0.0515 | 0.479 |  |  |  |
| **Pche-1::ire-1; ire-1(-):g.c.t vs. Pche-1::ire-1; ire-1(-):g.c.c** | **P<0.001** | **29.866** |  |  |  |
| **Fig.3 - supplement 1: Student's T-test** |  |  |  |  |  |
| Phsp-4::gfp; Pche-1::mcherry transgenic animals |  |  |  |  |  |
| **comparison** | **p-value** | **difference** |  |  |  |
| control RNAi vs. flp-6 RNAi | 0.0056 | 60.0884 |  |  |  |
| **Fig.3 - supplement 2: Student's T-test** |  |  |  |  |  |
| Pche-1::mcherry transgenic animals |  |  |  |  |  |
| **comparison** | **p-value** | **difference** |  |  |  |
| control RNAi vs. tfg-1 RNAi | 0.1258 | 0.0958 |  |  |  |
| **Fig.4A : two-way ANOVA followed by Tukey's post hoc analysis** |  |  |  |  |  |
| all animals treated with gld-1; ced-3 RNAi |  |  | degrees of freedom | p-value | F |
|  |  |  | 3(16) | <0.0001 | 28123.0402 |
| comparison | p-value | difference |  |  |  |
| AIB:Histamine + vs. AIB:Histamine - | P<0.001 | 0.7211 |  |  |  |
| RIA:Histamine + vs. RIA:Histamine - | P<0.001 | 0.9867 |  |  |  |
| AIA:Histamine + vs. AIA:Histamine - | 0.1669 | 0.2987 |  |  |  |
| AIY:Histamine + vs. AIY:Histamine - | P<0.001 | 35.4838 |  |  |  |
| **Fig.4B : One-way ANOVA followed by Tukey's post hoc analysis** |  |  |  |  |  |
| all animals treated with gld-1; ced-3 RNAi |  |  | degrees of freedom | p-value | F |
|  |  |  | 3(12) | <0.0001 | 3090.9606 |
| comparison | p-value | difference |  |  |  |
| control RNAi vs. AIY(-) | P<0.001 | 33.2808 |  |  |  |
| flp-6(-) vs. AIY(-) | 0.3167 | 0.7924 |  |  |  |
| flp-6(-); AIY(-) vs. AIY(-) | P<0.001 | 2.5845 |  |  |  |
| flp-6(-) vs. control RNAi | P<0.001 | 34.0732 |  |  |  |
| flp-6(-); AIY(-) vs. control RNAi | P<0.001 | 35.8652 |  |  |  |
| flp-6(-); AIY(-) vs. flp-6(-) | 0.0071 | 1.792 |  |  |  |
| **Fig.4C : One-way ANOVA followed by Tukey's post hoc analysis** |  |  |  |  |  |
| all animals treated with gld-1; ced-3 RNAi |  |  | degrees of freedom | p-value | F |
|  |  |  | 3(8) | <0.0001 | 6139.717 |
| comparison | p-value | difference |  |  |  |
| flp-6(-) vs. control(RNAi) | P<0.001 | 36.4714 |  |  |  |
| flp-6(-); mex-3(RNAi) vs. control(RNAi) | P<0.001 | 59.8202 |  |  |  |
| mex-3(RNAi) vs. control(RNAi) | P<0.001 | 25.3529 |  |  |  |
| flp-6(-); mex-3(RNAi) vs. flp-6(-) | P<0.001 | 23.3488 |  |  |  |
| mex-3(RNAi) vs. flp-6(-) | P<0.001 | 11.1184 |  |  |  |
| mex-3(RNAi) vs. flp-6(-); mex-3(RNAi) | P<0.001 | 34.4672 |  |  |  |
| **Fig.4D : One-way ANOVA followed by Tukey's post hoc analysis** |  |  |  |  |  |
| all animals treated with gld-1; ced-3 RNAi |  |  | degrees of freedom | p-value | F |
|  |  |  | 9(20) | <0.0001 | 13393.7324 |
| comparison | p-value | difference |  |  |  |
| wt vs. flp-6(-) | P<0.001 | 34.9672 |  |  |  |
| wt vs. flp-1(yn4) | 0.0019 | 1.0291 |  |  |  |
| wt vs. flp-1(yn2) | 0.3509 | 0.4985 |  |  |  |
| wt vs. flp-18(db99) | 0.0091 | 0.8851 |  |  |  |
| wt vs. flp-18(gk3063) | 1 | 0.0496 |  |  |  |
| wt vs. cha-1(-) | P<0.001 | 34.0733 |  |  |  |
| wt vs. Ptx-3::cha-1; cha-1(-) | 0.3801 | 0.4869 |  |  |  |
| wt vs. unc-17(-) | P<0.001 | 32.7788 |  |  |  |
| wt vs. Ptx-3::unc-17; unc-17(-) | 0.0043 | 0.9554 |  |  |  |
| Ptx-3::cha-1; cha-1(-) vs. cha-1(-) | P<0.001 | 34.5602 |  |  |  |
| Ptx-3::unc-17; unc-17(-) vs. unc-17(-) | P<0.001 | 33.7343 |  |  |  |
| **Fig.5A : One-way ANOVA followed by Tukey's post hoc analysis**<br>all animals treated with gld-1; ced-3 RNAi |  |  | **degrees of freedom** | **p-value** | **F** |
|  |  |  | 8(18) | <0.0001 | 12311.0006 |
| **comparison** | **p-value** | **difference** |  |  |  |
| wt vs. flp-6(ok3056) | P<0.001 | 46.3713 |  |  |  |
| wt vs. her-1(n695) | P<0.001 | 44.624 |  |  |  |
| wt vs. sem-4(ga82) | P<0.001 | 40.6063 |  |  |  |
| wt vs. mig-1(n1354) | P<0.001 | 43.1631 |  |  |  |
| wt vs. mig-19(k142) | P<0.001 | 42.1511 |  |  |  |
| wt vs. mig-20(k148) | P<0.001 | 42.1116 |  |  |  |
| wt vs. flp-6(-); sem-4(-) | P<0.001 | 36.302 |  |  |  |
| wt vs. flp-6(-); her-1(-) | P<0.001 | 34.1178 |  |  |  |
| flp-6(ok3056) vs. flp-6(-); sem-4(-) | P<0.001 | 10.0693 |  |  |  |
| sem-4(ga82) vs. flp-6(-); sem-4(-) | P<0.001 | 4.3043 |  |  |  |
| flp-6(ok3056) vs. flp-6(-); her-1(-) | P<0.001 | 12.2536 |  |  |  |
| her-1(n695) vs. flp-6(-); her-1(-) | P<0.001 | 10.5063 |  |  |  |
| **Fig.5B : One-way ANOVA followed by Tukey's post hoc analysis**<br>all animals treated with gld-1; ced-3 RNAi |  |  | **degrees of freedom** | **p-value** | **F** |
|  |  |  | 5(12) | <0.0001 | 297598.2607 |
| **comparison** | **p-value** | **difference** |  |  |  |
| wt vs. flp-6(-) | P<0.001 | 36.8307 |  |  |  |
| wt vs. nlp-3(-) | P<0.001 | 1.0325 |  |  |  |
| wt vs. nlp-8(-) | P<0.001 | 1.0054 |  |  |  |
| wt vs. nlp-15(-) | P<0.001 | 2.0277 |  |  |  |
| wt vs. flp-5(-) | P<0.001 | 2.9748 |  |  |  |
| **Fig.5C: One-way ANOVA followed by Tukey's post hoc analysis**<br>all animals treated with gld-1; ced-3 RNAi |  |  | **degrees of freedom** | **p-value** | **F** |
|  |  |  | 3(8) | <0.0001 | 2519.3729 |
| **comparison** | **p-value** | **difference** |  |  |  |
| wt vs. flp-6(-) | P<0.001 | 36.0799 |  |  |  |
| wt vs. tph-1(-) | P<0.001 | 35.2805 |  |  |  |
| wt vs. bas-1(-) | P<0.001 | 34.4234 |  |  |  |
| flp-6(-)-bas-1(-) | 0.0417 | 1.6565 |  |  |  |
| tph-1(-)-bas-1(-) | 0.3723 | 0.857 |  |  |  |
| tph-1(-)-flp-6(-) | 0.4259 | 0.7995 |  |  |  |
| **Fig.5D: One-way ANOVA followed by Tukey's post hoc analysis**<br>all animals treated with gld-1; ced-3 RNAi |  |  | **degrees of freedom** | **p-value** | **F** |
|  |  |  | 5(12) | <0.0001 | 45566.8396 |
| **comparison** | **p-value** | **difference** |  |  |  |
| wt + 20mM 5-HT vs. wt | P<0.001 | 1.0809 |  |  |  |
| tph-1(-) + 20mM 5-HT vs. tph-1(-) | P<0.001 | 29.5717 |  |  |  |
| flp-6(-) + 20mM 5-HT vs. flp-6(-) | P<0.001 | 34.0381 |  |  |  |
| **Fig.5E: One-way ANOVA followed by Tukey's post hoc analysis**<br>all animals treated with gld-1; ced-3 RNAi |  |  | **degrees of freedom** | **p-value** | **F** |
|  |  |  | 5(18) | <0.0001 | 14430.8317 |
| **comparison** | **p-value** | **difference** |  |  |  |
| wt-tph-1(-); genomic tph-1(+) | P<0.001 | 1.0173 |  |  |  |
| tph-1(-); genomic tph-1(+)-tph-1(-) in ADF | 0.9999 | 0.043 |  |  |  |
| tph-1(-); genomic tph-1(+)-tph-1(-) in ASG | 0.0722 | 0.5227 |  |  |  |
| tph-1(-); genomic tph-1(+)-tph-1(-) in HSN | P<0.001 | 28.1446 |  |  |  |
| wt-tph-1(-) | P<0.001 | 29.6247 |  |  |  |
| tph-1(-); genomic tph-1(+)-tph-1(-) | P<0.001 | 28.6074 |  |  |  |
| tph-1(-) in ADF-tph-1(-) | P<0.001 | 28.5644 |  |  |  |
| tph-1(-) in ASG-tph-1(-) | P<0.001 | 29.1301 |  |  |  |
| tph-1(-) in HSN-tph-1(-) | 0.1361 | 0.4628 |  |  |  |
| **Fig.5 - supplement 1 : One-way ANOVA followed by Tukey's post hoc analysis**<br>all animals treated with gld-1; ced-3 RNAi |  |  | **degrees of freedom** | **p-value** | **F** |
|  |  |  | 5(18) | <0.0001 | 7392.0341 |
| **comparison** | **p-value** | **difference** |  |  |  |
| wt-flp-6(-) | P<0.001 | 39.3757 |  |  |  |
| wt-mod-1(-) | 0.9973 | 0.1188 |  |  |  |
| wt-ser-1(-) | 1 | 0.0226 |  |  |  |
| wt-ser-4(-) | 0.4896 | 0.4779 |  |  |  |
| wt-ser-7(-) | 0.6629 | 0.4007 |  |  |  |

| Fig.6A : One-way ANOVA followed by Tukey's post hoc analysis |  |  |  |  |  |
| --- | --- | --- | --- | --- | --- |
| Punc-119::gfp; ced-3(n1286) animals treated with gld-1 RNAi |  |  | degrees of freedom | p-value | F |
|  |  |  | 5(12) | <0.0001 | 291.0161 |
| comparison | p-value | difference |  |  |  |
| control vs. che-1(p679) | P<0.001 | 26.0732 |  |  |  |
| control vs. flp-6(ok3056) | P<0.001 | 28.6601 |  |  |  |
| control vs. cha-1(p1152) | P<0.001 | 31.0352 |  |  |  |
| control vs. sem-4(n1378) | P<0.001 | 25.9739 |  |  |  |
| control vs. tph-1(mg280) | P<0.001 | 28.0861 |  |  |  |
| Difference represents the absolute difference between the means of the compared groups. |  |  |  |  |  |
| Note that while many of the comparisons are statistically significant, the extent of the observed changes vary substantially. |  |  |  |  |  |
| For experiments presenting percent of gonad area occupied by large nuclei, Differences beyond 20% are in bold. |  |  |  |  |  |

